## Supplementary Figures for "SMRT sequencing of the *Oryza rufipogon* genome reveals the genomic basis of rice adaptation"

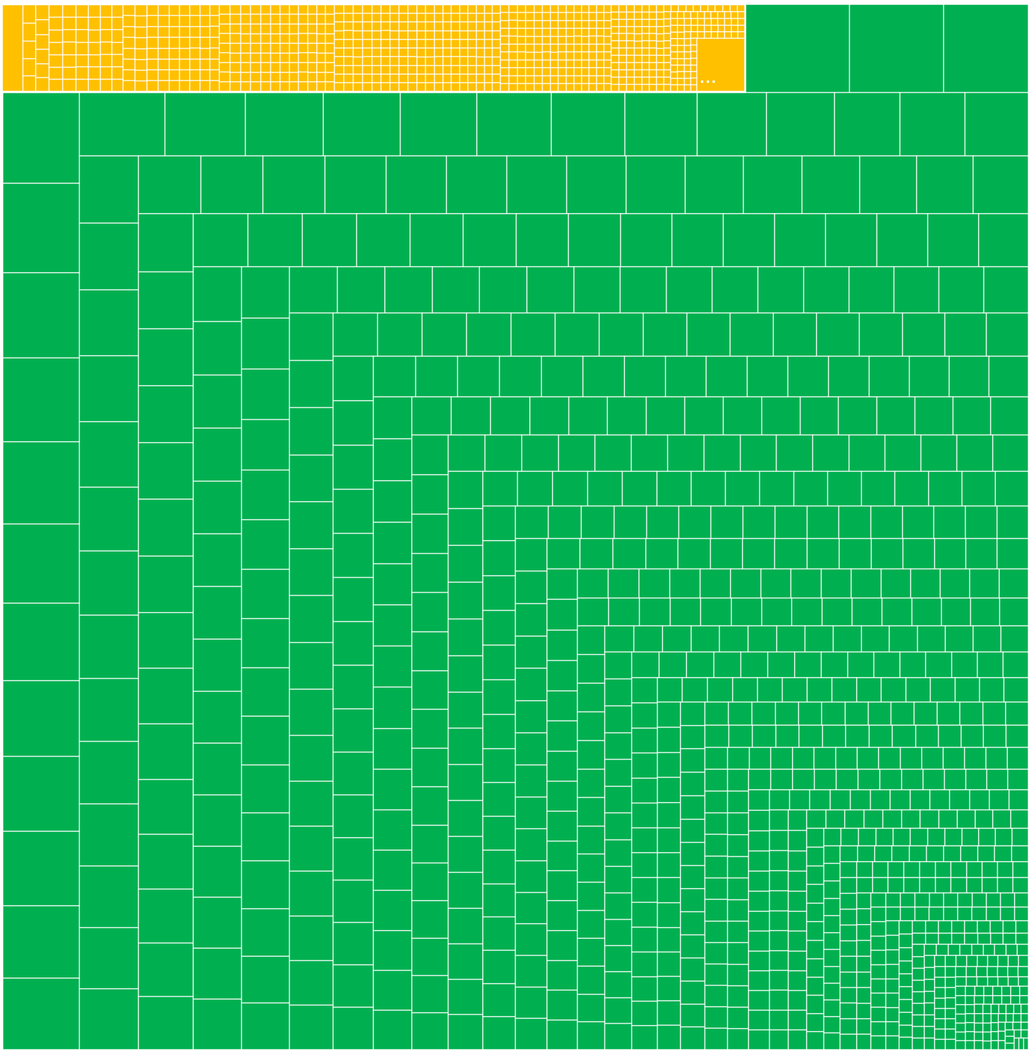

**Figure S1. Tree map of haplotigs (yellow) and p-contigs (green) scaled by length.**

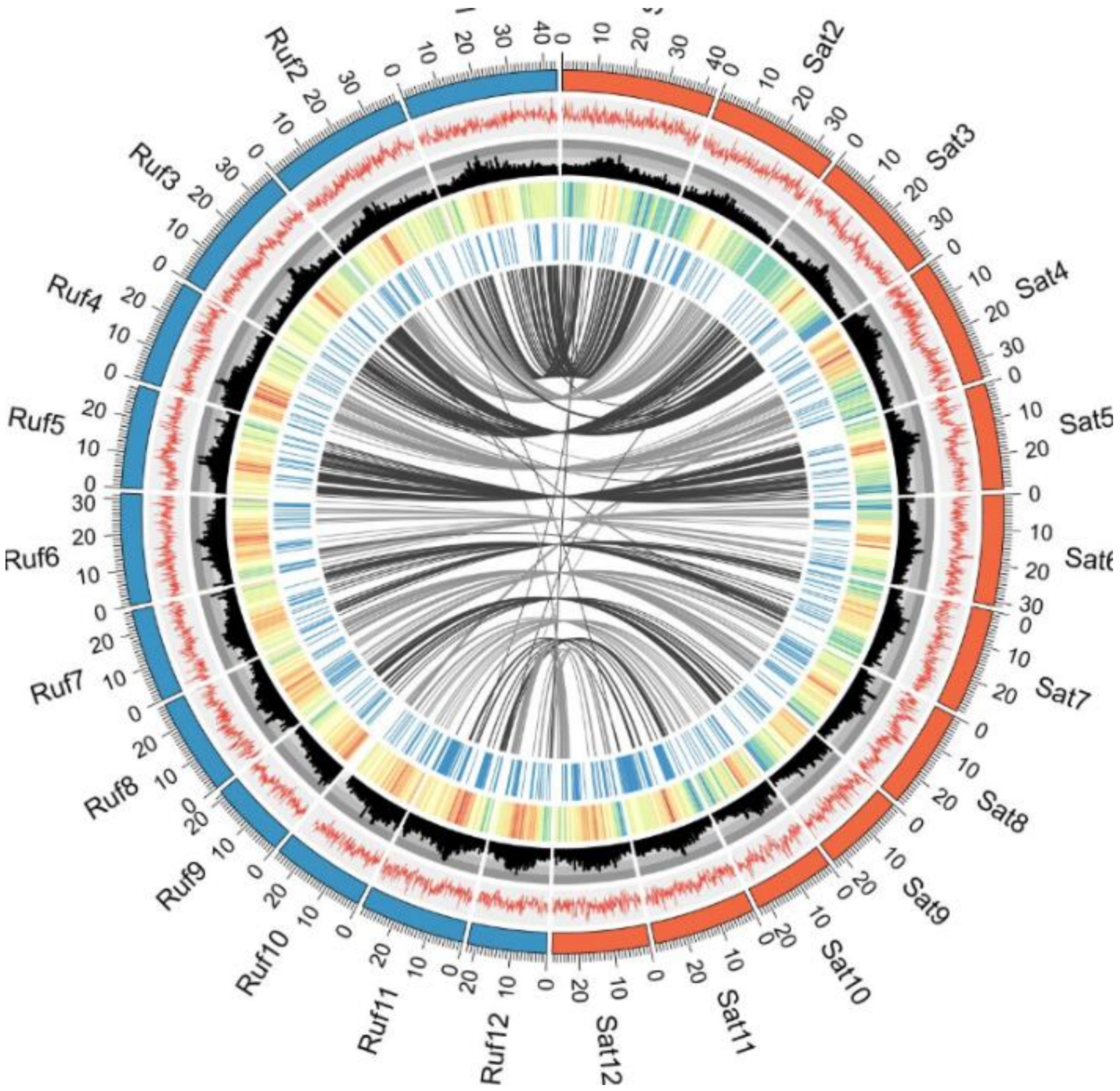


**Figure S2**. **Synteny comparisons and genome features of *O. rufipogon* and *O. sativa***. Tracks from outside to inside are the 12 chromosomes of *O. rufipogon* and *O. sativa*, GC content, transposable element (TE) density, gene density (density measured in 100-Kb sliding windows), and distribution of NBS genes. The syntenic blocks between *O. rufipogon* and *O. sativa* chromosomes are displayed with connecting lines in different colors.

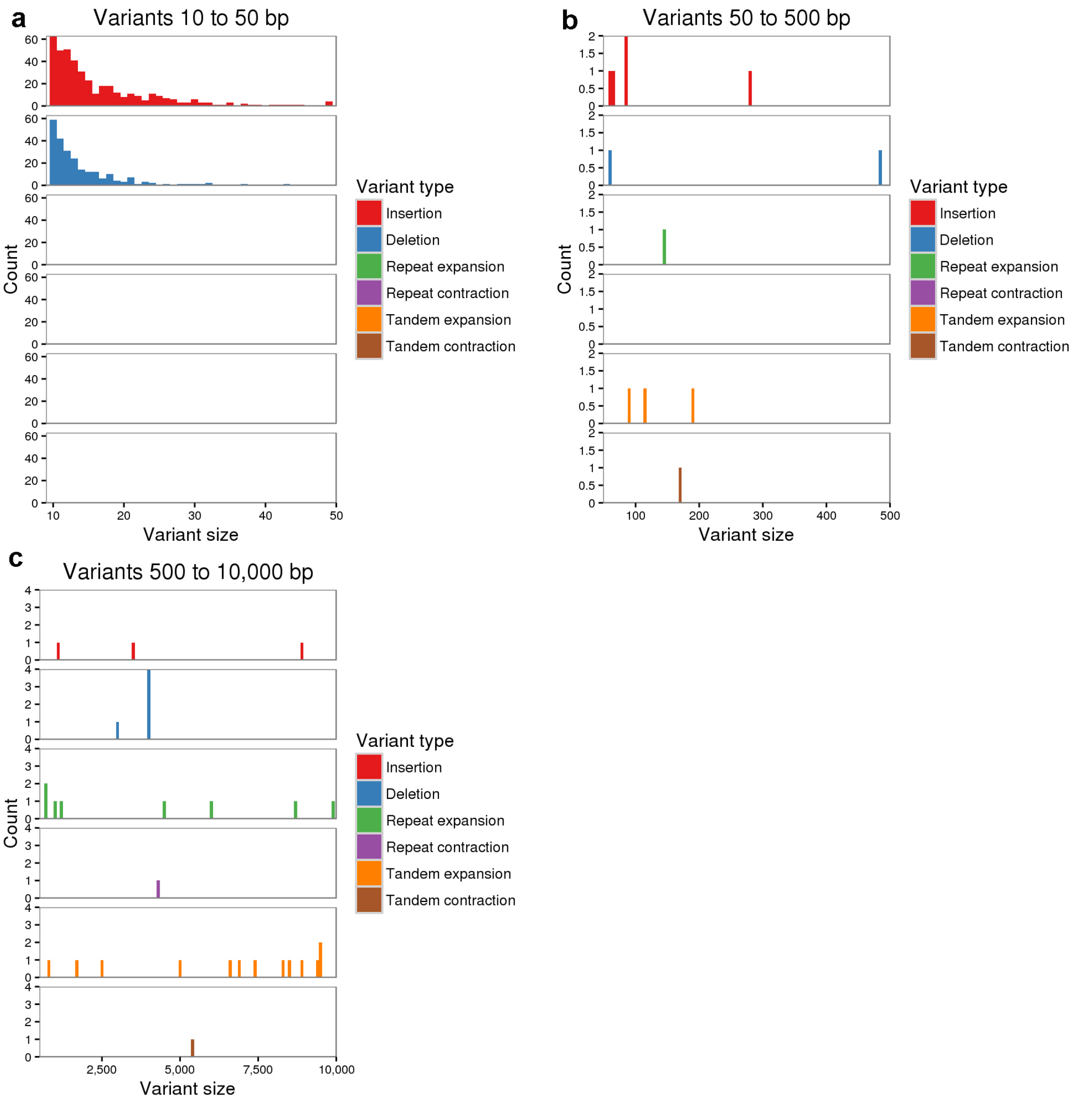


**Figure S3. Number of genomic variants in different lengths detected between haplotigs and p-contigs. (a)** Variants from 10 to 50 bp; **(b)** variants from 50 to 500 bp; **(c)** variants from 500 to 10,000 bp.

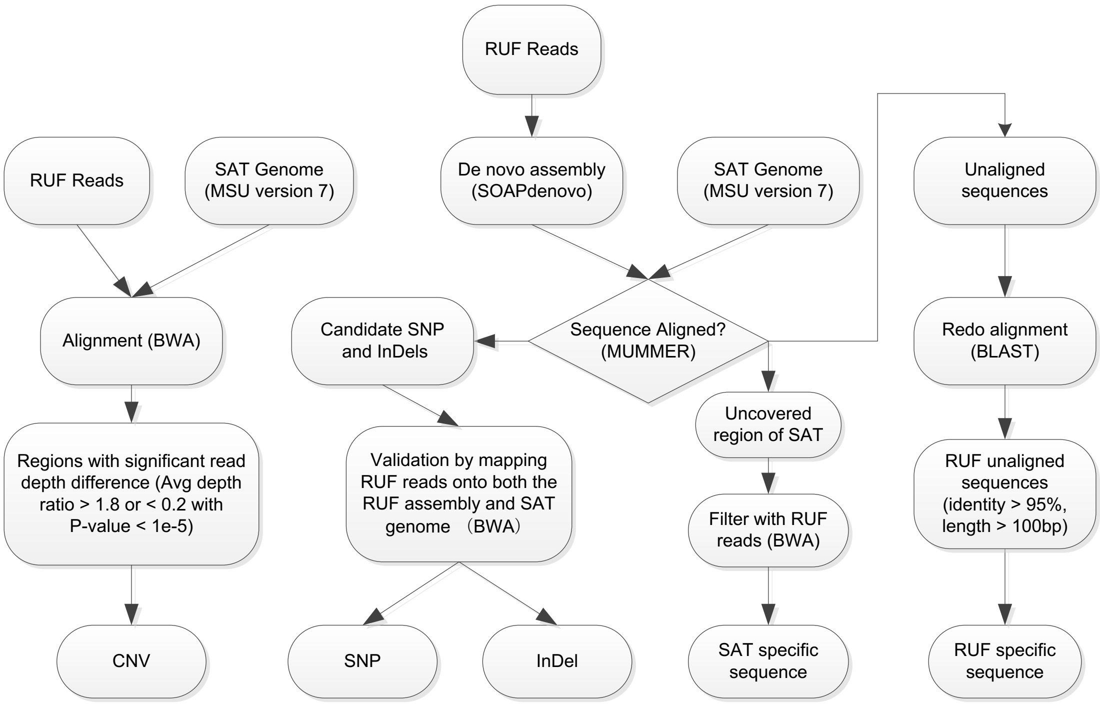


**Figure S4. Flowchart for the detection of genomic variation between RUF and the Nipponbare genome.** This pipeline was also applied to the comparison between the NIV and Nipponbare genomes.

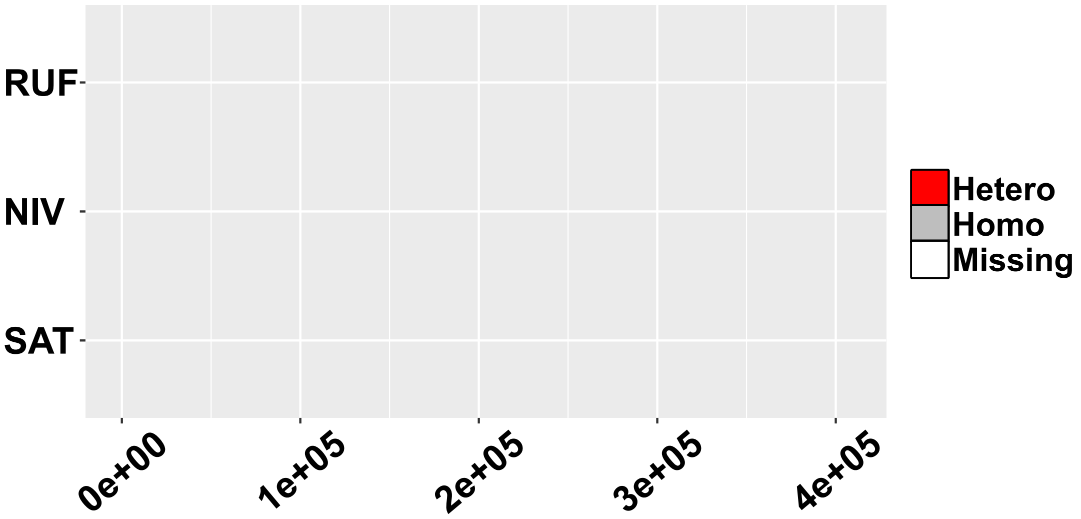


Chromosome 2

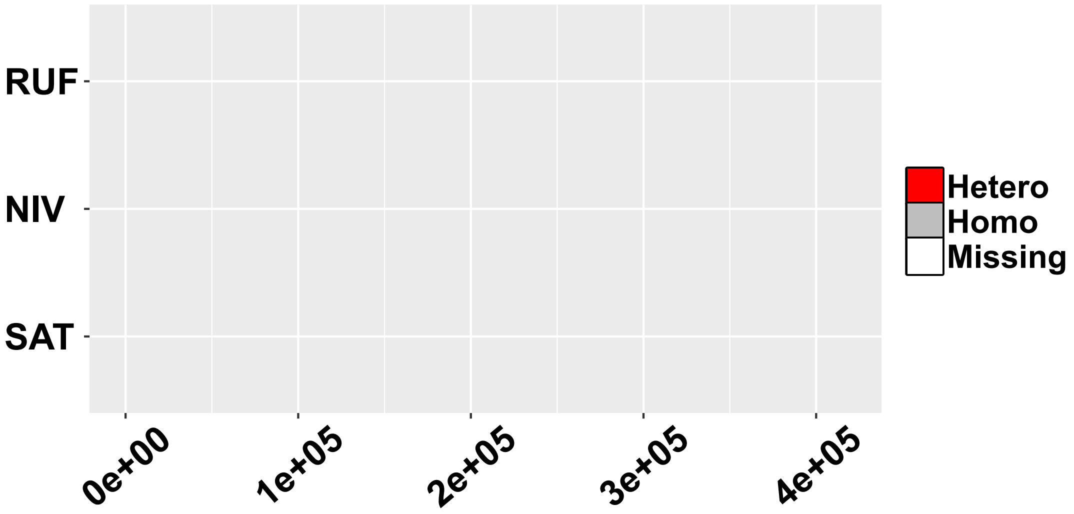


Chromosome 3

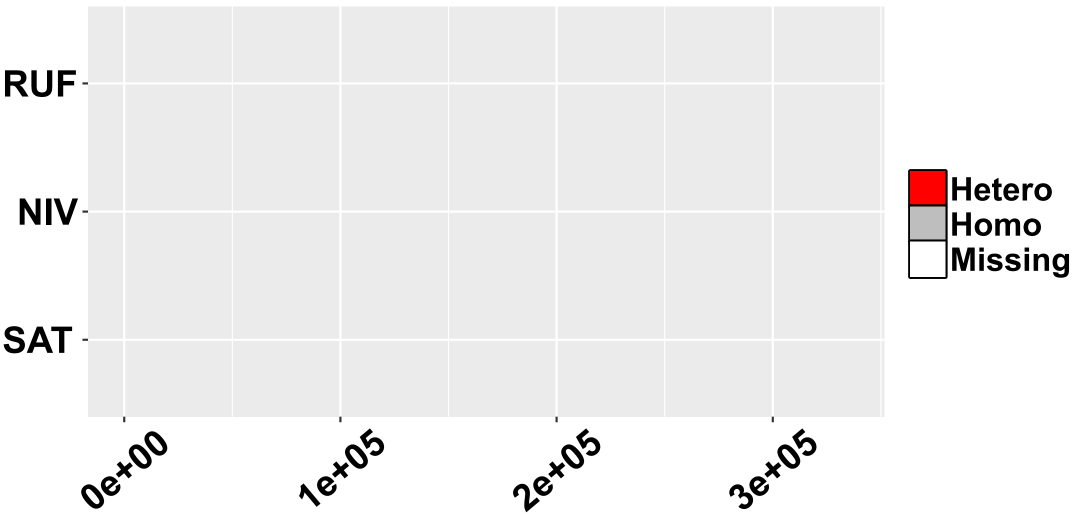


Chromosome 4

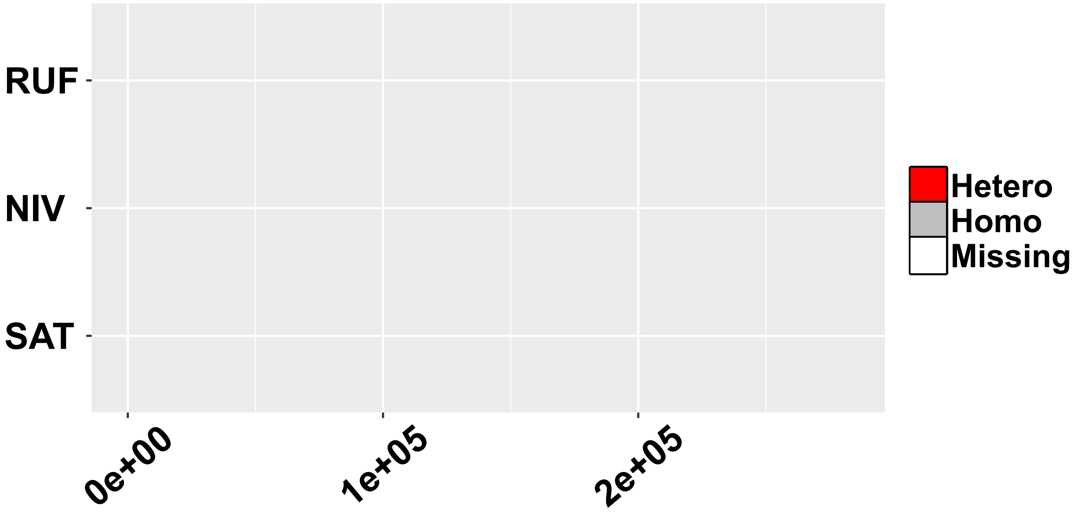


Chromosome 5

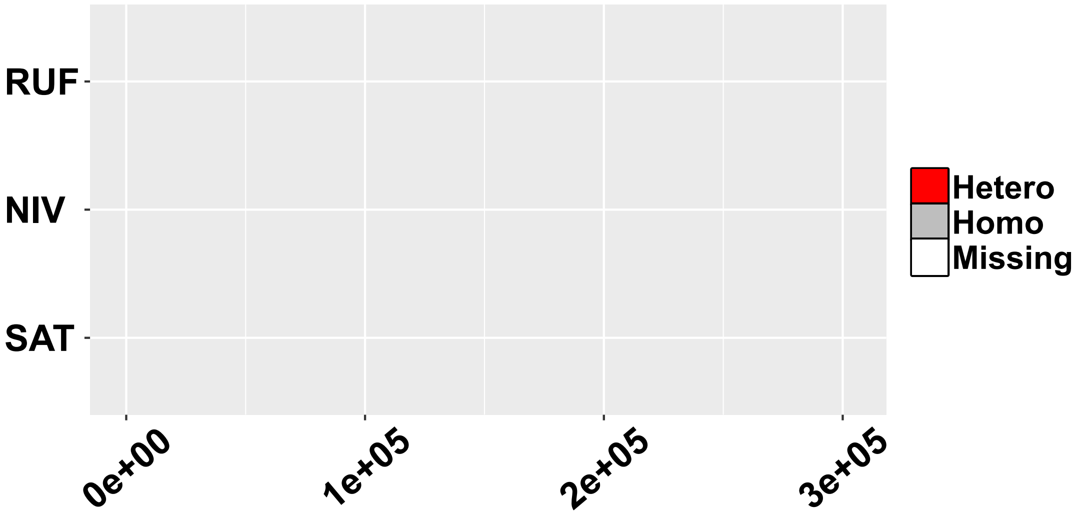


Chromosome 6

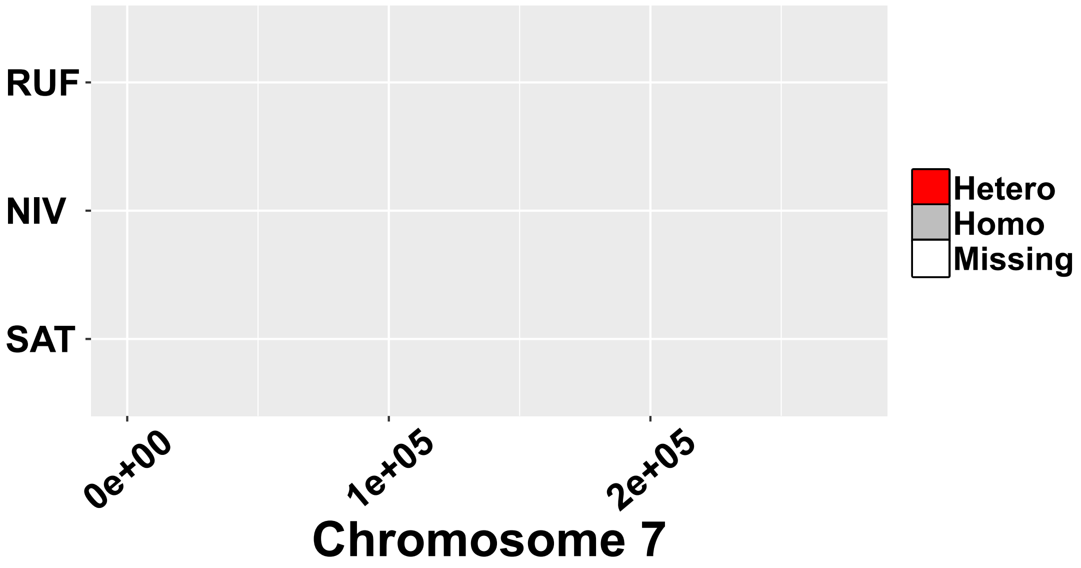
RUF
NIV
SAT
Hetero
Homo
Missing
0e+00
1e+05
2e+05
Chromosome 7

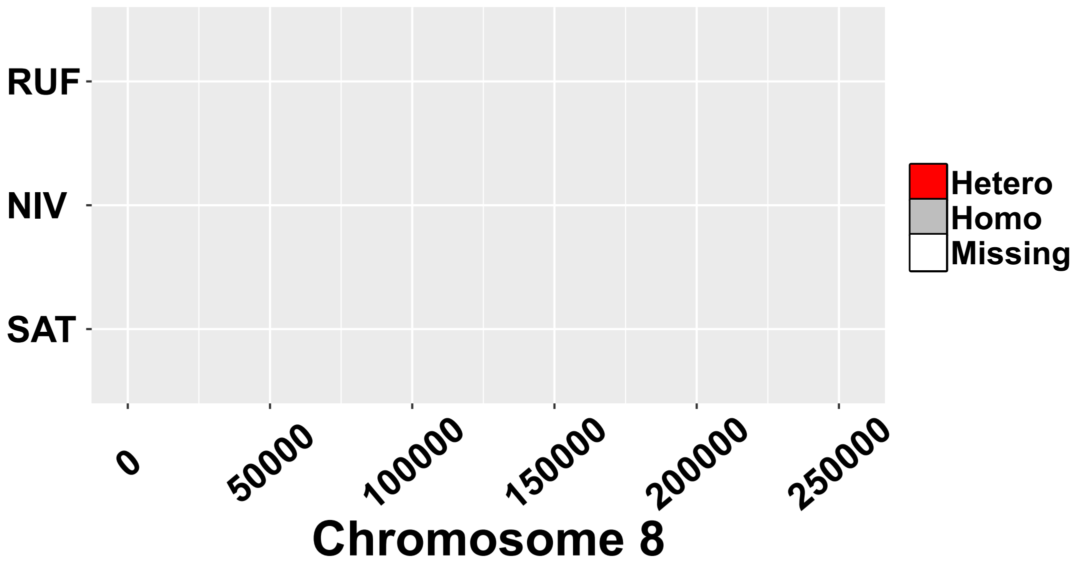
RUF
NIV
SAT
Hetero
Homo
Missing
0
50000
100000
150000
200000
250000
Chromosome 8

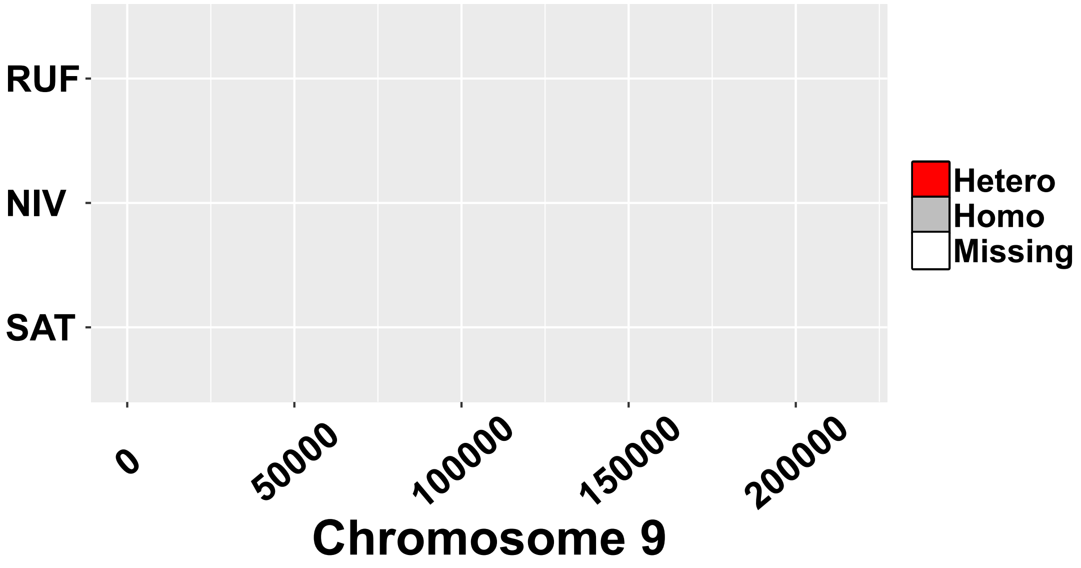
RUF
NIV
SAT
Hetero
Homo
Missing
0
50000
100000
150000
200000
Chromosome 9

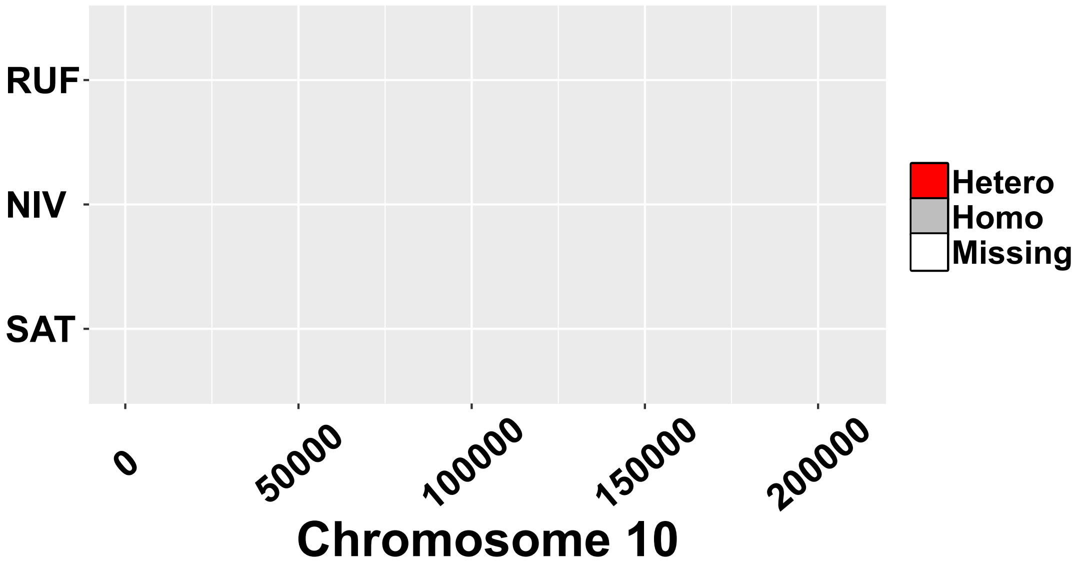
RUF
NIV
SAT
0
50000
100000
150000
200000
Chromosome 10
Hetero
Homo
Missing

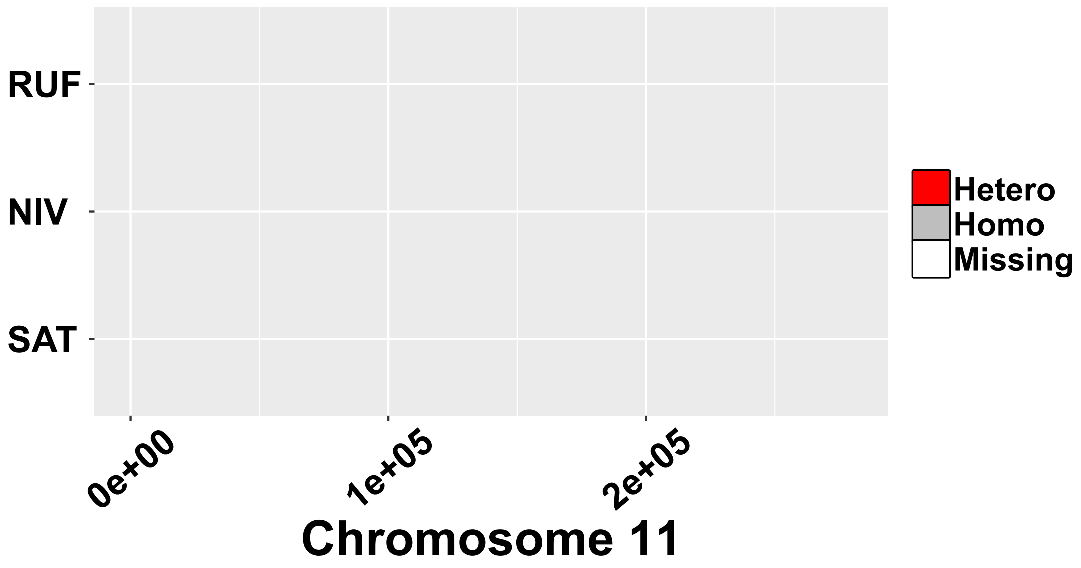
RUF
NIV
SAT
0e+00
1e+05
2e+05
Chromosome 11
Hetero
Homo
Missing

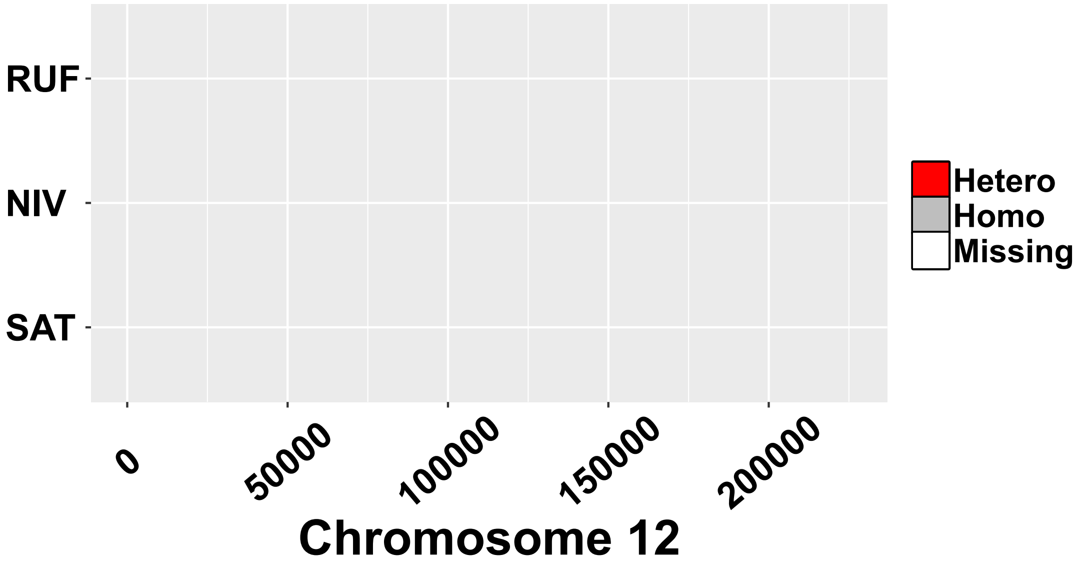
RUF
NIV
SAT
0
50000
100000
150000
200000
Chromosome 12
Hetero
Homo
Missing

**Figure S5. Patterns of single nucleotide polymorphisms along the 11 rice chromosomes among *O. rufipogon*, *O. nivara* and *O. sativa*.** Heterozygous and homozygous SNPs are shown with red and gray lines, respectively, while the unknown sites are indicated with white lines.

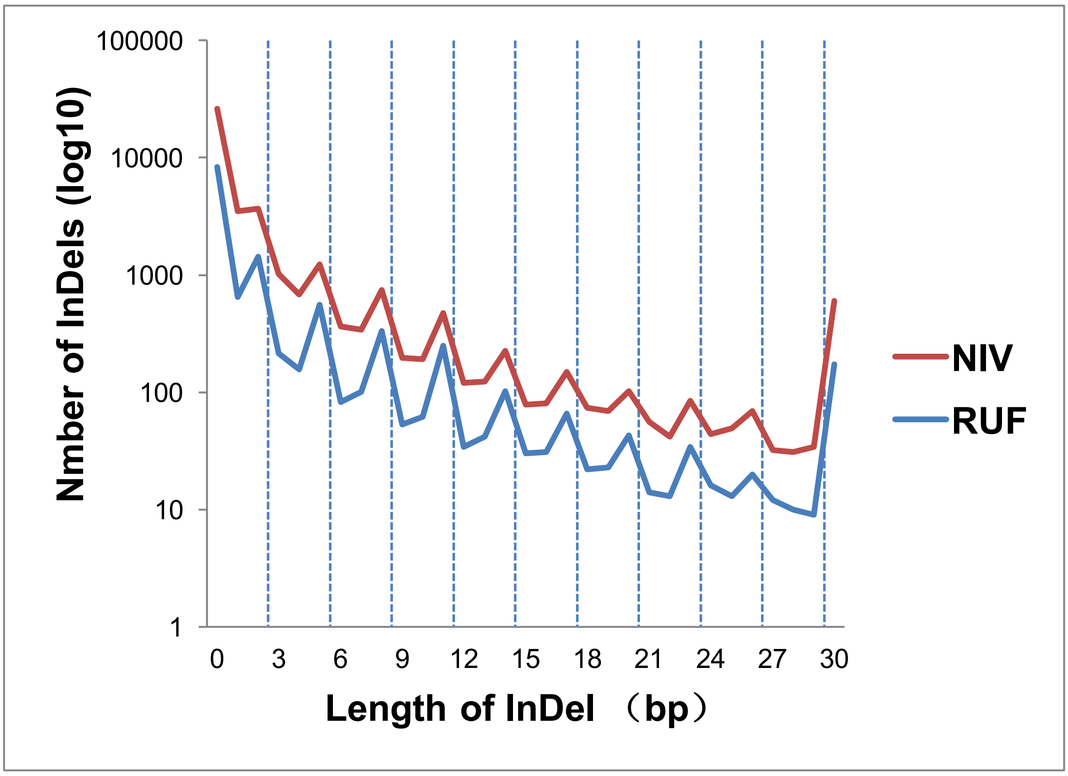


**Figure S6. Length distribution of InDels located within CDS of *O. rufipogon* and *O. nivara*.** The peaks at positions that are multiples of three are shown by dashed vertical lines.

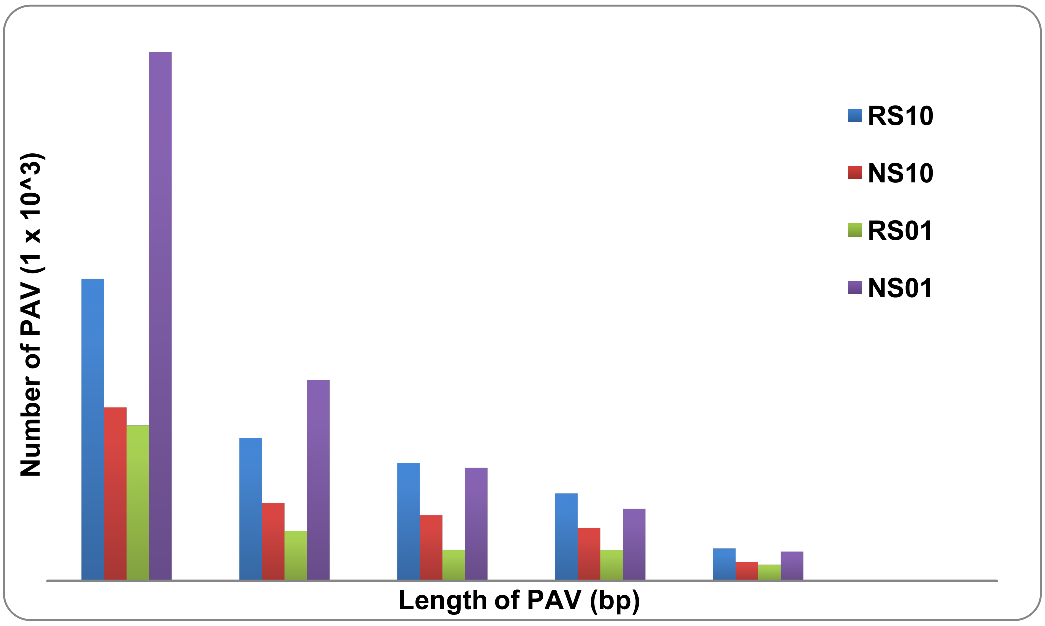


**Figure S7. Summary of PAVs among *O. rufipogon*, *O. nivara* and *O. sativa*.** R: *O. rufipogon* (RUF); S: *O. sativa* (SAT); N: *O. nivara* (NIV); 1: presence; 0: absence. We defined and assigned four types of PAV: RS10 (presence in RUF but absence in SAT), RS01 (presence in SAT but absence in RUF), NS10 (presence in NIV but absence in SAT) and NS01 (presence in SAT but absence in NIV).

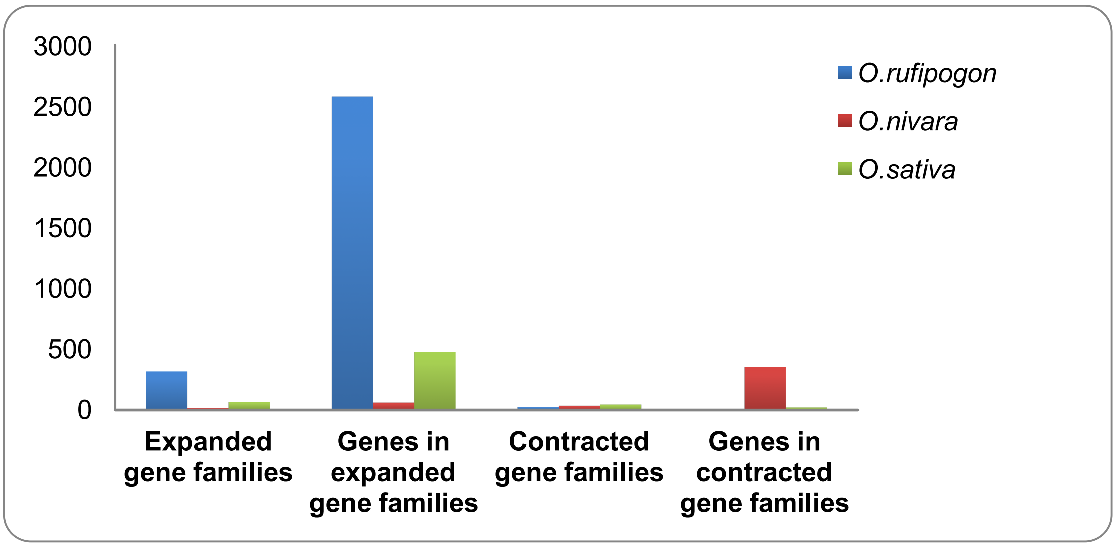


**Figure S8. Summary of gene families expanded or contracted significantly (*P* < 0.01) in terminal branches of *O. rufipogon*, *O. nivara* and *O. sativa*.**

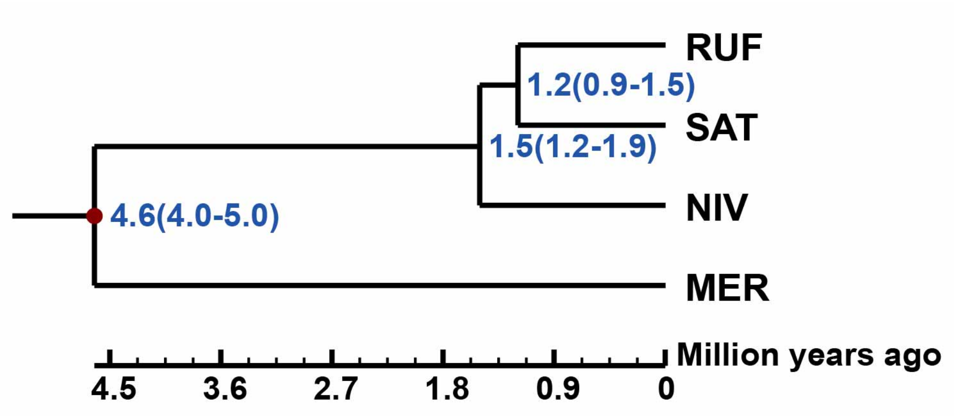


**Figure S9. Phylogenetic relationships and divergence times of *O. rufipogon* (RUF), *O. nivara* (NIV) and *O. sativa* (SAT) using *O. meridionalis* (MER) as outgroup.** The phylogeny was inferred from 10,206 high-confidence 1:1 orthologous gene families based on the maximum likelihood method under the GTR+ GAMMA model, and the divergence times were estimated using *mcmctree* program implemented in PAML. Note that all nodes are fully supported at 100%.

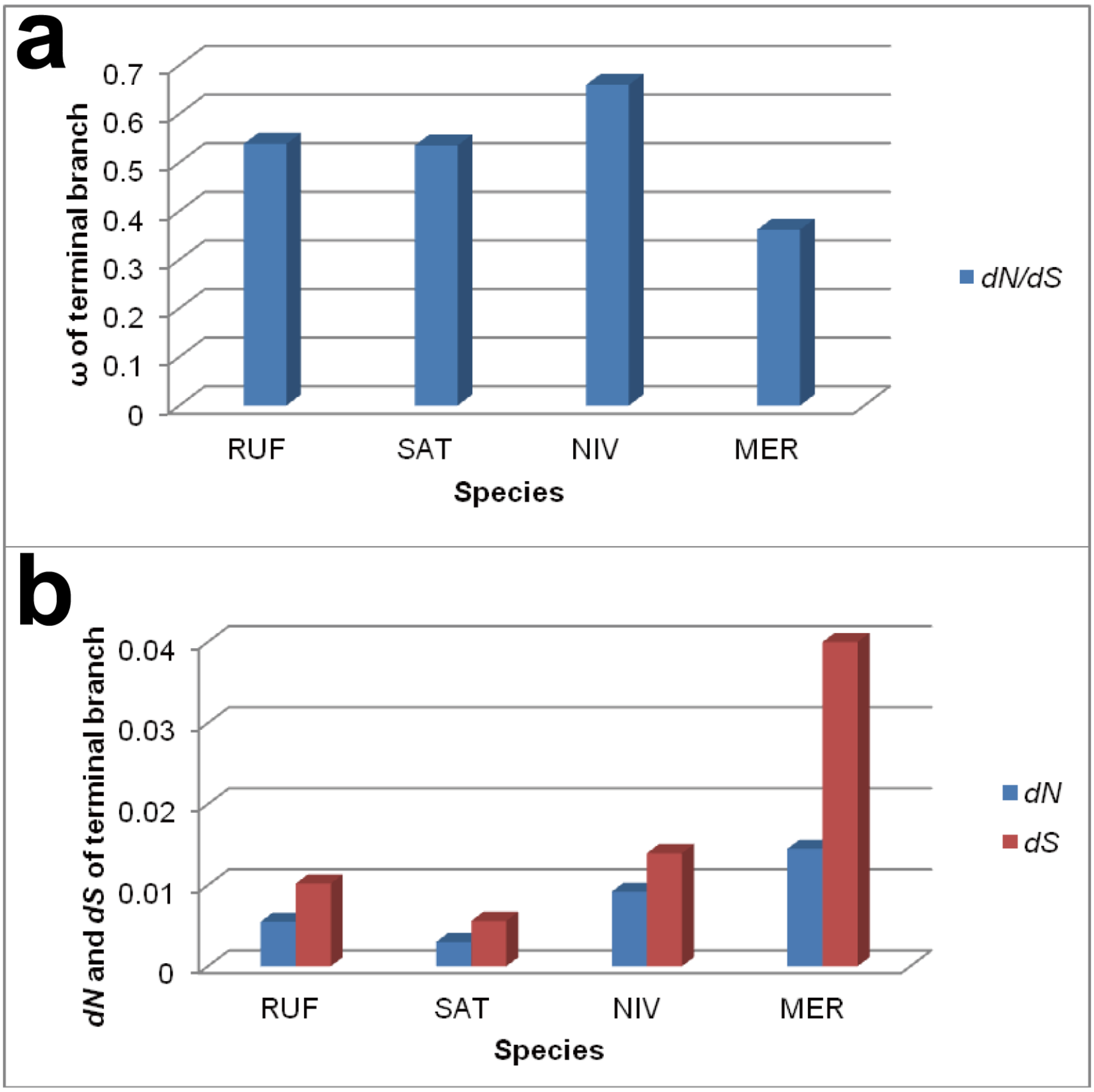


**Figure S10. Branch-specific *dN*, *dS* and ω values in each terminal branch.** Shown are values of (**a**) ω (*dN*/*dS*); (**b**) *dN* and *dS* for each species, which were estimated in branch model tests.
