## Supplementary Tables for "SMRT sequencing of the *Oryza rufipogon* genome reveals the genomic basis of rice adaptation"

**Table S1. Assembly statistics of the *O. rufipogon* genome.**

| | Assembler | | | |
|---|---|---|---|---|
| | **Falcon-Unzip** | **Falcon-Unzip+10×** | **Falcon-Unzip+10×+Hi-C** | **Falcon-Unzip+10×+Hi-C+PBJelly** |
| **Assembled Length (bp)** | 373,883,792 | 377,175,457 | 373,971,792 | 380,512,861 |
| **Scaffold N50 (bp)** | 710,332 | 2,214,952 | 29,494,264 | 30,197,710 |
| **Scaffold N90 (bp)** | 178,741 | 487,719 | 22,265,879 | 22,824,363 |
| **Contig N50 (bp)** | 710,332 | 710,332 | 710,332 | 1,096,430 |
| **Contig N90 (bp)** | 178,741 | 178,741 | 178,741 | 286,860 |
| **Scaffold Number** | 1,404 | 843 | 524 | 524 |
| **Contig Number** | 1,404 | 1,404 | 1,404 | 1,077 |
| **Longest Scaffold (bp)** | 4,290,542 | 7,550,243 | 43,695,925 | 44,260,698 |
| **Longest Contig (bp)** | 4,290,542 | 4,290,542 | 4,290,542 | 4,409,078 |

**Table S2. Assembly results of p-contigs and haplotigs of the *O. rufipogon* genome..**

| Sequences | Assembly length (Mb) | No. contigs | N50 length (Kb) | N90 length (Kb) | Max contig length (Kb) |
|---|---|---|---|---|---|
| **P-contigs** | 373.88 | 1,404 | 710.33 | 178.74 | 4,290.54 |
| **Haplotigs** | 23.85 | 843 | 29.47 | 18.51 | 653.91 |

**Table S3. Scaffold length constitutions of *O. rufipogon*.**

| Scaffold Length | Number | Scaffold Length (bp) | Average Length (bp) | Percentage (%) |
|---|---|---|---|---|
| **>1 Kb** | 826 | 377,165,817 | 456,617 | 100.00 |
| **>10 Kb** | 600 | 375,914,405 | 626,524 | 99.67 |
| **>50 Kb** | 331 | 370,046,179 | 1,117,964 | 98.11 |
| **>100 Kb** | 290 | 367,168,813 | 1,266,099 | 97.35 |
| **>200 Kb** | 252 | 361,676,028 | 1,435,222 | 95.89 |
| **>300 Kb** | 230 | 356,507,333 | 1,550,031 | 94.52 |
| **>500 Kb** | 184 | 337,877,604 | 1,836,291 | 89.58 |
| **>800 Kb** | 141 | 311,420,982 | 2,208,659 | 82.57 |
| **>1 Mb** | 122 | 294,593,993 | 2,414,705 | 78.11 |
| **>2 Mb** | 59 | 204,848,868 | 3,472,015 | 54.31 |
| **>3 Mb** | 29 | 132,571,336 | 4,571,425 | 35.15 |

**Table S4. Assembly statistics for the *O. rufipogon* genome sequences.**

| Chromosome ID | Chromosome Length (RUF) (bp) | Chromosome length (Nipponbare) (bp) |
|---|---|---|
| 1 | 43,695,925 | 43,270,923 |
| 2 | 39,958,125 | 35,937,250 |
| 3 | 37,088,850 | 36,413,819 |
| 4 | 29,494,264 | 35,502,694 |
| 5 | 28,993,391 | 29,958,434 |
| 6 | 31,585,856 | 31,248,787 |
| 7 | 26,272,893 | 29,697,621 |
| 8 | 27,299,189 | 28,443,022 |
| 9 | 21,773,623 | 23,012,720 |
| 10 | 26,444,676 | 23,207,287 |
| 11 | 29,583,747 | 29,021,106 |
| 12 | 22,265,879 | 27,531,856 |
| Unmapped | 9,515,374 | 0 |
| Total | 373,971,792 | 373,245,519 |

**Table S5. Statistics for large variants detected between haplotigs and p-contigs.**

| Variant Type | Count | Total length (bp) |
|---|---|---|
| **Insertion** | 429 | 21,114 |
| **Deletion** | 247 | 22,989 |
| **Tandem_expansion** | 16 | 85,296 |
| **Tandem_contraction** | 2 | 5,617 |
| **Repeat_expansion** | 9 | 32,690 |
| **Repeat_contraction** | 1 | 4,301 |
| **Total** | 704 | 172,007 |

**Table S6. Quality assessment of the *O. rufipogon* genome assembly using reads mapping, DNA, protein and EST datasets.**

| | Total | Aligned | Percentage (%) |
|---|---|---|---|
| **Reads mapping** | | | |
| PE reads | 175,474,936 | 163,187,767 | 93.00 |
| **Sequences available in public databases** | | | |
| DNA * | 20,667 | 17,968 | 86.94 |
| Protein ** | 3,762 | 2,424 | 64.43 |
| **Transcripts of *O. rufipogon*** | | | |
| EST * | 105,654 | 75,374 | 71.34 |
| **BUSCOs from Embryophyta lineage*** ** | | | |
| Complete | 1,440 | 1,402 | 97.36 |
| Duplicated | 1,440 | 17 | 1.18 |
| Fragmented | 1,440 | 11 | 0.76 |
| Missing | 1,440 | 27 | 1.88 |

*Aligned using GMAP (version 2014-10-22), and the hits with identity ≥ 90% && coverage ≥ 90% are retained.

**Aligned using genBlastA (version 1.0.1), and the hits with identity ≥ 80% && coverage ≥ 90% are retained.

***The 1,440 BUSCO conserved genes used were collected from Embryophyta lineage.

**Table S7. Summary of gene prediction for *O. rufipogon.***

| Type | | Values |
|---|---|---|
| | Total number of predicted genes (#) | 34,830 |
| | Total number of gene models (#) | 34,830 |
| | Average gene length (bp) | 2,921 |
| **Gene Features** | Average CDS length (bp) | 1,125 |
| | Average exons per gene | 4.5 |
| | Average exon length (bp) | 248 |
| | Average intron length (bp) | 507 |

**Table S8. Validation of gene models of *O. rufipogon* by using homologous proteins and RNA-Seq datasets.**

| Evidence | Resources | Number | % Percent |
|---|---|---|---|
| Total predicted genes | This study | 34,589 | 100 |
| Homologous protein supported* | *O. sativa* | 24,711 | 71.44 |
| RNA-Seq supported** | This study | 21,515 | 62.20 |
| Homology protein or RNA-Seq supported | | 29,123 | 84.20 |

Note: Blast with e-value < $1e^{-5}$ was used for searching against database;

* Identity >= 30% and Coverage >= 80%;

** Identity >= 90% and Coverage >= 50%

**Table S9. Summary of non-coding RNA genes in the *O. rufipogon*.**

| Type | Number | Average Length (bp) | Total Length (bp) | % of Genome |
|---|---|---|---|---|
| tRNA | 637 | 75 | 47,862 | 0.0127 |
| rRNA (8S) | 976 | 116 | 113,091 | 0.0300 |
| rRNA (18S) | 56 | 1,814 | 101,573 | 0.0269 |
| rRNA (28S) | 53 | 3,933 | 208,438 | 0.0553 |
| SnoRNA | 442 | 120 | 53,026 | 0.0141 |
| snRNA | 117 | 143 | 16,713 | 0.0044 |
| miRNA | 245 | 140 | 34,236 | 0.0091 |

**Table S10. Statistics of repeat sequence in the *O. rufipogon* genome.**

| | Length (bp) | Percentage (%) |
|---|---|---|
| **Transposable Elements** | 166467134 | 44.14 |
| **DNA transposons** | 56904501 | 15.09 |
| ***En-Spm*** | 13033370 | 3.46 |
| ***Harbinger*** | 5602726 | 1.49 |
| ***MuDR*** | 15643183 | 4.15 |
| ***TcMar-Stowaway*** | 3271049 | 0.87 |
| ***Tourist*** | 14,711 | 0.00 |
| ***hAT*** | 3900448 | 1.03 |
| ***Helitron*** | 11628837 | 3.08 |
| **Other** | 3810177 | 1.01 |
| **Retrotransposon** | 102758700 | 27.24 |
| **Non-LTR Retrotransposon** | 5174466 | 1.37 |
| **LINE** | 4498862 | 1.19 |
| **SINE** | 675604 | 0.18 |
| **LTR Retrotransposon** | 97584234 | 25.87 |
| ***Copia*** | 11212454 | 2.97 |
| ***Gypsy*** | 58639158 | 15.55 |
| **Other*** | 27732622 | 7.35 |
| **Other Repeats** | 6803933 | 1.80 |
| **Satellite** | 1168 | 0.00 |
| **Simple repeats** | 4799297 | 1.27 |
| **Unknown** | 2003468 | 0.53 |

* Non-autonomous LTR retrotransposons.

**Table S11. Occurrence of simple sequence repeats (SSRs) in the *O. rufipogon* genome.**

| Repeat type | Number | Proportion (%) | Total length (Kb) | Average length (bp) |
|---|---|---|---|---|
| Mononucleotide | 16790 | 7.67 | 238.82 | 14 |
| Dinucleotide | 39838 | 18.19 | 964.59 | 24 |
| Trinucleotide | 84516 | 38.60 | 1190.43 | 14 |
| Tetranucleotide | 49983 | 22.83 | 659.63 | 13 |
| Pentanucleotide | 16967 | 7.75 | 271.29 | 16 |
| Hexanucleotide | 10873 | 4.97 | 208.58 | 19 |
| **Total** | 218967 | 100.00 | 3533.34 | 16 |

**Table S12. Assembly statistics of the *O. rufipogon*, *O.sativa* and *O. nivara* genomes.**

| | ***O.sativa*** | ***O. nivara*** | ***O. rufipogon*** |
|---|---|---|---|
| **Sequencing technology** | Sanger | Illumina | PacBio |
| **Completeness (BUSCO) (%)** | 98.2 | 95.4 | 97.36 |
| **Contig N50 (bp)** | 7,711,345 | 13,868 | 1,096,430 |
| **Scaffold N50 (bp)** | 29,958,434 | 506,802 | 30,197,710 |
| **Total length (bp)** | 374,471,240 | 379,501,719 | 380,512,861 |
| **Protein No.** | 39,045 | 41,490 | 34,830 |

**Table S13. Summary of the pan-genome of *O. rufipogon*, *O. sativa* and *O. nivara*.**

| | Sequence length (bp) |
|---|---|
| Shared among RUF, NIV and SAT | 317,729,226 |
| Shared between RUF and NIV | 349,534,715 |
| Shared between RUF and SAT | 342,883,315 |
| Shared between NIV and SAT | 336,486,012 |
| RUF-specific | 98,013,406 |
| NIV-specific | 11,210,218 |
| SAT-specific | 12,831,139 |
| Pan-genome | 515,500,353 |

Note: *O. sativa* is abbreviated as SAT, *O. rufipogon* is abbreviated as RUF and *O. nivara* is abbreviated as NIV.

**Table S14. Enrichment of core and dispensable genes in various GO categories.**

| | GO term | Frequency in core genes | Frequency in dispensable genes | *P*-value | FDR |
|---|---|---|---|---|---|
| **Core genes** | | | | | |
| BP | Biological regulation | 0.0988 | 0.0747 | 2.19E-06 | 2.63E-05 |
| BP | Regulation of biological process | 0.0970 | 0.0739 | 5.05E-06 | 3.03E-05 |
| CP | Cell | 0.1165 | 0.0923 | 1.13E-05 | 4.51E-05 |
| CP | Cell part | 0.1165 | 0.0923 | 1.13E-05 | 3.38E-05 |
| BP | Cellular component organization or biogenesis | 0.0261 | 0.0166 | 2.57E-04 | 6.18E-04 |
| BP | Localization | 0.0859 | 0.0699 | 7.08E-04 | 1.42E-03 |
| BP | Cellular process | 0.3798 | 0.3527 | 1.21E-03 | 2.08E-03 |
| BP | Single-organism process | 0.2467 | 0.2230 | 1.31E-03 | 1.97E-03 |
| CP | Organelle | 0.0784 | 0.0646 | 1.95E-03 | 2.60E-03 |
| MF | Nucleic acid binding transcription factora ctivity | 0.0353 | 0.0267 | 4.23E-03 | 5.07E-03 |
| CP | Membrane part | 0.0675 | 0.0596 | 4.11E-02 | 4.48E-02 |
| MF | Protein binding transcription factor activity | 0.0036 | 0.0020 | 4.86E-02 | 4.86E-02 |
| **Dispensable genes** | | | | | |
| BP | Reproductive process | 0.0026 | 0.0123 | 9.26E-11 | 9.26E-10 |
| BP | Multi-organism process | 0.0031 | 0.0127 | 3.69E-10 | 1.85E-09 |
| BP | Multicellular organismal process | 0.0054 | 0.0150 | 3.91E-08 | 1.30E-07 |
| MF | Nutrient reservoir activity | 0.0025 | 0.0073 | 4.98E-05 | 1.24E-04 |
| MF | Structural molecule activity | 0.0136 | 0.0216 | 4.01E-04 | 8.03E-04 |
| MF | Catalytic activity | 0.4345 | 0.4578 | 5.62E-03 | 9.37E-03 |
| BP | Growth | 0.0004 | 0.0016 | 2.09E-02 | 2.98E-02 |
| MF | Enzyme regulator activity | 0.0090 | 0.0123 | 4.27E-02 | 5.34E-02 |
| MF | Electron carrier activity | 0.0087 | 0.0119 | 4.53E-02 | 5.03E-02 |
| MF | Metallochaperone activity | 0.0000 | 0.0004 | 4.96E-02 | 4.96E-02 |

**Table S15. Summary of genomic variation among *O. sativa*, *O. rufipogon* and *O. nivara*.**

**A. Combined method of genome comparison and reads mapping analysis**

| **Species** | **Genomic Variation** | | | |
|---|---|---|---|---|
| | Number of SNP | Number of InDel | Number of SV | Number of CNV |
| RUF | 4,997,466 | 817,238 | 16,163 | 5,017 |
| NIV | 3,794,980 | 779,252 | 19,914 | 7,766 |
| SAT | 23,584 | 2,764 | 1,879 | 2,034 |

Note: Numbers of SNP and InDel were obtained from whole genome comparisons and reads mapping analysis; SV and CNV calling was only based on reads mapping analysis.

**B. Genome comparison method**

| **Species** | **Genomic Variation** | |
|---|---|---|
| | Number of SNP | Number of InDel |
| RUF | 2,887,048 | 551,018 |
| NIV | 3,093,758 | 619,797 |

Note: SNP and InDel calling are based on whole genome alignments.

**C. Reads mapping method**

| **Species** | **Genomic Variation** | | | |
|---|---|---|---|---|
| | Number of SNP | Number of InDel | Number of SV | Number of CNV |
| RUF | 4,065,179 | 403,544 | 16,163 | 5,017 |
| NIV | 2,841,519 | 319,771 | 19,914 | 7,766 |
| SAT | 23,584 | 2,764 | 1,879 | 2,034 |

Note: *O. sativa* is abbreviated as SAT, *O. rufipogon* is abbreviated as RUF and *O. nivara* is abbreviated as NIV.

**Table S16. Annotation of genomic variation among *O. sativa*, *O. rufipogon* and *O. nivara*.**

**A. SNP annotation**

| **Species** | **Number of SNP** | | | |
|---|---|---|---|---|
| | Number of Synonymous Sites | Number of Non-synonymous Sites | Number of Stop codon Gain | Number of Stop codon Loss |
| RUF | 339,831 | 446,309 | 17,124 | 2,218 |
| NIV | 238,642 | 349,519 | 14,083 | 1,730 |

**B. InDel annotation**

| **Species** | **Number of InDel** | | | |
|---|---|---|---|---|
| | Number of Stop codon Loss | Number of Stop codon Gain | Number of FrameShift | Number of Non-FrameShift |
| RUF | 109 | 920 | 25,139 | 14,614 |
| NIV | 88 | 1,242 | 41,038 | 13,568 |

**C. CNV annotation**

| **Species** | **Number of CNV** | |
|---|---|---|
| | Number of Deletion | Number of Duplication |
| RUF | 3,355 | 1,662 |
| NIV | 3,562 | 4,204 |
| SAT | 1,806 | 228 |

Note: Deletion and duplication refer that copy number fewer and more than in the reference genome, respectively.

**D. SV annotation**

| **Species** | **SV** | | | | | | |
|---|---|---|---|---|---|---|---|
| | Number of ICT1 | Number of Deletion | Number of Insertion | Number of Inversion | Number of ITC2 | Number of Exon | |
| | | | | | | Number of FrameShift | Number of Non-FrameShift |
| RUF | 52 | 12,035 | 0 | 465 | 621 | 4,539 | 266 |
| NIV | 185 | 13,655 | 2,524 | 384 | 431 | 4,660 | 554 |
| SAT | 3 | 47 | 1,357 | 26 | 20 | 228 | 91 |

Note: ICT1 refers to Intra-chromosome translocation; ITX2 refers to Inter-chromosome translocation. SVs that occur within the exon regions were classified them into two types, frameshift and non-frameshift. *O. sativa* is abbreviated as SAT, *O. rufipogon* is abbreviated as RUF and *O. nivara* is abbreviated as NIV.

**Table S17. Shared and sample-specific genes affected by CNVs in the *O. rufipogon* and *O. nivara* genomes.**

| | Number of shared genes | |
|---|---|---|
| | N=2 | N=1 |
| Loss | 88 | 940 |
| Gain | 145 | 6,940 |
| Loss/Gain | 86 | - |
| Total | 319 | 7,880 |

**Table S21. Numbers of MADS-box genes affected by CNVs and total MADS-box genes from whole genome of *O. sativa*.**

| MADS-box type | Affected by CNVs | Whole genome |
|---|---|---|
| M-α | 4 | 13 |
| M-β | 2 | 9 |
| M-γ | 0 | 10 |
| **Type I (subtotal)** | 6 | 32 |
| MIKC$^C$ | 15 | 37 |
| MIKC* | 2 | 5 |
| **Type II (subtotal)** | 17 | 42 |
| **Total** | 23 | 74 |

**Table S22. The 23 MADS-box genes affected by CNVs in *O. sativa*.**

| Gene ID | Physical location | Type | #Wild species | MADS-box type |
|---|---|---|---|---|
| LOC_Os01g10504 | 5559548-5568844 | CNV-gain | 1 | MIKCc |
| LOC_Os01g66290 | 38500880-38505127 | CNV-gain | 1 | MIKCc |
| LOC_Os01g67890 | 39459599-39461050 | CNV-gain | 1 | M-beta |
| LOC_Os01g69850 | 40344329-40364584 | CNV-gain | 1 | MIKC* |
| LOC_Os02g07430 | 3833129-3837135 | CNV-gain | 1 | MIKCc |
| LOC_Os02g36924 | 22294657-22301808 | CNV-gain | 1 | MIKCc |
| LOC_Os02g52340 | 32038902-32045130 | CNV-gain | 1 | MIKCc |
| LOC_Os03g11614 | 6052750-6061369 | CNV-gain | 1 | MIKCc |
| LOC_Os04g23910 | 13672710-13675884 | CNV-gain | 1 | MIKCc |
| LOC_Os04g31804 | 19042703-19058654 | CNV-gain | 1 | M-alpha |
| LOC_Os04g52410 | 31143083-31144886 | CNV-gain | 1 | MIKCc |
| LOC_Os05g23780 | 13656346-13657002 | CNV-gain | 1 | M-alpha |
| LOC_Os06g06750 | 3162801-3169415 | CNV-gain | 1 | MIKCc |
| LOC_Os06g45650 | 27637555-27641578 | CNV-gain | 1 | MIKCc |
| LOC_Os07g04170 | 1781625-1785426 | CNV-gain | 1 | M-beta |
| LOC_Os07g41370 | 24788476-24793884 | CNV-gain | 1 | MIKCc |
| LOC_Os08g02070 | 679358-681739 | CNV-gain | 1 | MIKCc |
| LOC_Os08g20440 | 12272625-12277708 | CNV-gain | 1 | M-alpha |
| LOC_Os08g33488 | 20897215-20906881 | CNV-gain | 1 | MIKCc |
| LOC_Os08g41950 | 26507180-26512261 | CNV-gain | 1 | MIKCc |
| LOC_Os11g43740 | 26414394-26418442 | CNV-gain | 1 | MIKC* |
| LOC_Os12g10540 | 5584593-5590285 | CNV-gain | 1 | MIKCc |
| LOC_Os12g21850 | 12303478-12304062 | CNV-gain | 1 | M-alpha |

**Table S23. Numbers of flowering-related genes affected by CNVs and total flowering-related genes from whole genome of *O. sativa*.**

| Flowering-related genes | Affected by CNVs | Whole genome |
|---|---|---|
| *CAB* | 4 | 14 |
| *CCA* | 5 | 18 |
| *CDF* | 0 | 5 |
| *CO* | 1 | 7 |
| *COP* | 1 | 1 |
| *CRY* | 1 | 6 |
| *ELF* | 0 | 3 |
| *FKF* | 0 | 1 |
| *FT* | 4 | 18 |
| *GI* | 1 | 1 |
| *LFY* | 0 | 1 |
| *LHY* | 0 | 1 |
| *Lux* | 6 | 29 |
| *PHOT* | 6 | 27 |
| *PHY* | 2 | 3 |
| *PIF_PIL* | 0 | 2 |
| *PRR* | 3 | 19 |
| *SOC* | 17 | 53 |
| *TOC* | 0 | 1 |
| *ZTL* | 2 | 4 |
| **Total** | **53** | **210** |

* Flowering-related genes in rice were detected using BLASTP (Identity ≥ 30% && Coverage ≥ 70%) based on the homologous protein sequences from *Arabidopsis thaliana*.

**Table S24. The 53 flowering-related genes affected by CNVs in *O. sativa*.**

| Gene ID | Physical location | Type | #Wild species | Symbol |
|---|---|---|---|---|
| LOC_Os01g08700 | 4329152-4338486 | CNV-gain | 1 | *GI* |
| LOC_Os01g10504 | 5559548-5568844 | CNV-gain | 1 | *SOC* |
| LOC_Os01g18800 | 10622795-10627552 | CNV-gain | 1 | *PHOT* |
| LOC_Os01g64360 | 37358204-37359596 | CNV-loss | 1 | *CCA* |
| LOC_Os01g64970 | 37710241-37715296 | CNV-gain | 1 | *PHOT* |
| LOC_Os01g66290 | 38500880-38505127 | CNV-gain | 1 | *SOC* |
| LOC_Os02g04640 | 2075496-2079149 | CNV-gain | 1 | *Lux* |
| LOC_Os02g07430 | 3833129-3837135 | CNV-gain | 1 | *SOC* |
| LOC_Os02g10390 | 5468367-5470496 | CNV-gain | 1 | *CAB* |
| LOC_Os02g36924 | 22294657-22301808 | CNV-gain | 1 | *SOC* |
| LOC_Os02g39090 | 23609300-23610643 | CNV-loss | 1 | *PHOT* |
| LOC_Os02g45670 | 27776453-27781443 | CNV-gain | 1 | *CCA* |
| LOC_Os02g47190 | 28808920-28811976 | CNV-gain | 1 | *Lux* |
| LOC_Os02g52340 | 32038902-32045130 | CNV-gain | 1 | *SOC* |
| LOC_Os02g53140 | 32528037-32533583 | CNV-gain | 1 | *COP* |
| LOC_Os02g58350 | 35689247-35690390 | CNV-gain | 1 | *PRR* |
| LOC_Os03g11614 | 6052750-6061369 | CNV-gain | 1 | *SOC* |
| LOC_Os03g14840 | 8087161-8088952 | CNV-loss | 1 | *PHOT* |
| LOC_Os03g17570 | 9759479-9768690 | CNV-gain | 1 | *PRR* |
| LOC_Os03g19590 | 11020091-11028228 | CNV-gain | 1 | *PHY* |
| LOC_Os03g51030 | 29168142-29176112 | CNV-gain | 1 | *PHY* |
| LOC_Os04g23910 | 13672710-13675884 | CNV-gain | 1 | *SOC* |
| LOC_Os04g41130 | 24391002-24395755 | CNV-gain | 1 | *FT* |
| LOC_Os04g52410 | 31143083-31144886 | CNV-gain | 1 | *SOC* |
| LOC_Os05g23780 | 13656346-13657002 | CNV-gain | 1 | *SOC* |
| LOC_Os05g50340 | 28845445-28846050 | CNV-gain | 1 | *CCA* |
| LOC_Os06g01670 | 398591-403494 | CNV-gain | 1 | *CCA* |
| LOC_Os06g06750 | 3162801-3169415 | CNV-gain | 1 | *SOC* |
| LOC_Os06g16370 | 9336359-9338643 | CNV-gain | 1 | *CO* |

| | | | | |
|---|---|---|---|---|
| LOC_Os06g45100 | 27275439-27278872 | CNV-gain | 1 | *CRY* |
| LOC_Os06g45650 | 27637555-27641578 | CNV-gain | 1 | *SOC* |
| LOC_Os06g45840 | 27734722-27740731 | CNV-gain | 1 | *CCA* |
| LOC_Os06g47890 | 28969488-28974576 | CNV-gain | 1 | *ZTL* |
| LOC_Os07g02800 | 1046017-1048052 | CNV-gain | 1 | *Lux* |
| LOC_Os07g25710 | 14753753-14759375 | CNV-gain | 1 | *Lux* |
| LOC_Os07g37550 | 22487421-22488766 | CNV-gain | 1 | *CAB* |
| LOC_Os07g41370 | 24788476-24793884 | CNV-gain | 1 | *SOC* |
| LOC_Os07g48596 | 29091952-29094140 | CNV-gain | 1 | *Lux* |
| LOC_Os07g49460 | 29616705-29629223 | CNV-gain | 1 | *PRR* |
| LOC_Os08g02070 | 679358-681739 | CNV-gain | 1 | *SOC* |
| LOC_Os08g06370 | 3530967-3535437 | CNV-gain | 1 | *Lux* |
| LOC_Os08g13360 | 7946528-7951938 | CNV-gain | 1 | *ZTL* |
| LOC_Os08g33488 | 20897215-20906881 | CNV-gain | 1 | *SOC* |
| LOC_Os08g33820 | 21171214-21172680 | CNV-gain | 1 | *CAB* |
| LOC_Os08g41950 | 26507180-26512261 | CNV-gain | 1 | *SOC* |
| LOC_Os09g33850 | 19981179-19986301 | CNV-gain | 1 | *FT* |
| LOC_Os11g05320 | 2356476-2361247 | CNV-gain | 1 | *PHOT* |
| LOC_Os11g13890 | 7659682-7662281 | CNV-gain | 1 | *CAB* |
| LOC_Os11g18870 | 10732922-10734798 | CNV-gain | 1 | *FT* |
| LOC_Os12g05590 | 2565213-2566562 | CNV-gain | 1 | *FT* |
| LOC_Os12g10540 | 5584593-5590285 | CNV-gain | 1 | *SOC* |
| LOC_Os12g21850 | 12303478-12304062 | CNV-gain | 1 | *SOC* |
| LOC_Os12g29580 | 17628062-17632426 | CNV-gain | 1 | *PHOT* |

**Table S25. Rice flower development pathway-associated genes affected by CNVs.**

| GeneID | Physical location | Type | Function | Symbol |
|---|---|---|---|---|
| LOC_Os01g07790 | 3737893-3740685 | CNV-gain | Polygalacturonase, putative, expressed | *PG* |
| LOC_Os01g23740 | 13353629-13357510 | CNV-gain | OsPDIL2-2 protein disulfide isomerase PDIL2-2, expressed | *PDIL* |
| LOC_Os02g34850 | 20899808-20907175 | CNV-gain | Histone-lysine N-methyltransferase ASHH2, putative, expressed | *ASHH2* |
| LOC_Os03g18200 | 10204819-10210306 | CNV-gain | Heat shock protein DnaJ, putative, expressed | *TMS1* |
| LOC_Os04g34450 | 20862474-20872470 | CNV-gain | Expressed protein | *SEC5* |
| LOC_Os04g49450 | 29500951-29503775 | CNV-gain | MYB family transcription factor, putative, expressed | *LHY* |
| LOC_Os05g05310 | 2611683-2617251 | CNV-gain | Fibronectin type III domain containing protein, expressed | *VIL1* |
| LOC_Os07g06970 | 3429443-3434747 | CNV-gain | HEN1, putative, expressed | *HEN1* |
| LOC_Os09g13610 | 7914083-7925405 | CNV-gain | PFT1, putative, expressed | *PFT1* |
| LOC_Os10g02770 | 1092789-1096966 | CNV-gain | Glycosyl hydrolases family 16, putative, expressed | *XTH* |
| LOC_Os10g27470 | 14487133-14493136 | CNV-gain | KH domain containing protein, putative, expressed | *PEP* |

**Table S26. Number of *R*-genes affected by CNVs and total *R*-genes from whole genome of *O. sativa*.**

| Type | Located in CNV region | Whole genome |
|---|---|---|
| CC-NBS | 8 | 30 |
| CC-NBS-LRR | 34 | 73 |
| NBS-LRR | 96 | 322 |
| TIR-NBS | 1 | 1 |
| NBS only | 57 | 186 |
| Total | 196 | 612 |

**Table S27. The 196 *R*-genes affected by CNVs in *O. sativa*.** * Number of wild species.

| Gene ID | Physical location | Type | Wild rice* | *R*-gene Type |
|---|---|---|---|---|
| LOC_Os01g02250 | 689792-694013 | CNV-loss | 1 | NBS-LRR |
| LOC_Os01g02280 | 707340-714322 | CNV-loss | 1 | NBS-LRR |
| LOC_Os01g05600 | 2669952-2672924 | CNV-gain | 1 | NBS-LRR |
| LOC_Os01g05620 | 2682019-2684988 | CNV-gain | 1 | NBS-LRR |
| LOC_Os01g07870 | 3804528-3811069 | CNV-gain | 1 | NBS |
| LOC_Os01g33684 | 18531902-18540820 | CNV-loss | 1 | CC-NBS-LRR |
| LOC_Os01g33810 | 18613744-18616560 | CNV-gain | 1 | CC-NBS-LRR |
| LOC_Os01g39990 | 22552874-22554531 | CNV-gain | 1 | CC-NBS |
| LOC_Os01g50080 | 28774281-28780365 | CNV-gain | 1 | NBS |
| LOC_Os01g50100 | 28786109-28793300 | CNV-gain | 1 | NBS |
| LOC_Os01g55530 | 31992087-31996466 | CNV-gain | 1 | NBS-TIR |
| LOC_Os01g57270 | 33091621-33096363 | CNV-loss | 1 | NBS-LRR |
| LOC_Os01g57280 | 33098072-33104550 | CNV-loss | 1 | NBS-LRR |
| LOC_Os01g57310 | 33116117-33124371 | CNV-loss | 2 | NBS-LRR |
| LOC_Os01g70080 | 40556007-40561361 | CNV-loss | 1 | NBS-LRR |
| LOC_Os01g72390 | 41986078-41988856 | CNV-loss | 1 | NBS-LRR |
| LOC_Os01g72410 | 41997451-42000861 | CNV-loss | 1 | NBS-LRR |
| LOC_Os02g02640 | 967172-974497 | CNV-loss | 1 | CC-NBS-LRR |
| LOC_Os02g02660 | 978126-987135 | CNV-loss | 1 | NBS-LRR |
| LOC_Os02g02670 | 988442-995506 | CNV-loss | 1 | CC-NBS-LRR |
| LOC_Os02g09790 | 5048656-5052503 | CNV-gain | 2 | CC-NBS-LRR |
| LOC_Os02g10900 | 5785295-5788769 | CNV-gain | 1 | NBS |
| LOC_Os02g18080 | 10503049-10508541 | CNV-gain | 1 | NBS-LRR |
| LOC_Os02g18140 | 10535635-10539699 | CNV-gain | 1 | NBS-LRR |
| LOC_Os02g18180 | 10552429-10558888 | CNV-gain | 1 | NBS |
| LOC_Os02g18510 | 10776284-10780758 | CNV-gain | 2 | NBS-LRR |
| LOC_Os02g18670 | 10880504-10886278 | CNV-gain | 2 | NBS |
| LOC_Os02g18700 | 10908431-10916390 | CNV-loss | 1 | NBS |
| LOC_Os02g19890 | 11701937-11708395 | CNV-gain | 1 | NBS-LRR |

| | | | | |
|---|---|---|---|---|
| LOC_Os02g27540 | 16311925-16315043 | CNV-gain | 1 | NBS-LRR |
| LOC_Os02g41760 | 25103334-25105616 | CNV-gain | 1 | NBS |
| LOC_Os03g05730 | 2852293-2856682 | CNV-gain | 1 | NBS |
| LOC_Os03g10900 | 5596830-5599784 | CNV-loss | 1 | NBS-LRR |
| LOC_Os03g38250 | 21233600-21237936 | CNV-gain | 1 | NBS-LRR |
| LOC_Os03g38330 | 21268409-21272412 | CNV-gain | 1 | NBS-LRR |
| LOC_Os03g48320 | 27517616-27521492 | CNV-gain | 1 | NBS-LRR |
| LOC_Os03g63200 | 35710304-35715776 | CNV-loss | 2 | NBS-LRR |
| LOC_Os04g02860 | 1113475-1118619 | CNV-gain | 1 | CC-NBS-LRR |
| LOC_Os04g13210 | 7301573-7312295 | CNV-gain | 1 | NBS |
| LOC_Os04g13220 | 7317232-7325788 | CNV-gain | 1 | NBS |
| LOC_Os04g21110 | 11869125-11887514 | CNV-gain | 1 | NBS |
| LOC_Os04g21890 | 12396117-12398915 | CNV-gain | 1 | NBS-LRR |
| LOC_Os04g30610 | 18279293-18281498 | CNV-loss | 1 | NBS |
| LOC_Os04g30660 | 18318639-18322459 | CNV-loss | 1 | NBS-LRR |
| LOC_Os04g30690 | 18340866-18344512 | CNV-loss | 1 | NBS |
| LOC_Os04g30930 | 18490353-18494159 | CNV-gain | 1 | NBS-LRR |
| LOC_Os04g40290 | 23955470-23959975 | CNV-gain | 1 | NBS |
| LOC_Os04g46300 | 27430842-27434629 | CNV-gain | 1 | NBS-LRR |
| LOC_Os04g52690 | 31364043-31369487 | CNV-gain | 1 | NBS |
| LOC_Os04g52970 | 31553065-31558406 | CNV-gain | 1 | NBS |
| LOC_Os04g53496 | 31854429-31862286 | CNV-gain | 1 | NBS-LRR |
| LOC_Os05g15040 | 8526403-8535211 | CNV-gain | 1 | NBS-LRR |
| LOC_Os05g30220 | 17512760-17517933 | CNV-gain | 1 | NBS-LRR |
| LOC_Os05g32370 | 18872317-18875899 | CNV-gain | 1 | NBS |
| LOC_Os05g34220 | 20258661-20262474 | CNV-gain | 1 | NBS-LRR |
| LOC_Os05g34230 | 20263661-20266481 | CNV-gain | 1 | NBS-LRR |
| LOC_Os05g40150 | 23580975-23585400 | CNV-gain | 1 | NBS-LRR |
| LOC_Os05g40160 | 23586768-23590400 | CNV-gain | 1 | NBS-LRR |
| LOC_Os05g45690 | 26472204-26473772 | CNV-gain | 1 | NBS |
| LOC_Os05g47490 | 27203619-27210015 | CNV-gain | 1 | NBS |
| LOC_Os05g47500 | 27211913-27217567 | CNV-gain | 1 | NBS |
| LOC_Os05g50790 | 29132047-29135068 | CNV-loss | 1 | NBS |

| | | | | |
|---|---|---|---|---|
| LOC_Os06g01980 | 550916-555929 | CNV-gain | 1 | NBS |
| LOC_Os06g06380 | 2973521-2979496 | CNV-gain | 1 | NBS-LRR |
| LOC_Os06g06390 | 2980288-2984308 | CNV-gain | 1 | NBS-LRR |
| LOC_Os06g06400 | 2985094-2990867 | CNV-gain | 1 | NBS-LRR |
| LOC_Os06g06440 | 3001105-3009250 | CNV-gain | 1 | NBS |
| LOC_Os06g06850 | 3234519-3239156 | CNV-gain | 1 | NBS-LRR |
| LOC_Os06g06860 | 3241818-3246312 | CNV-gain | 1 | NBS-LRR |
| LOC_Os06g15750 | 8935092-8939018 | CNV-gain | 1 | CC-NBS-LRR |
| LOC_Os06g16450 | 9418321-9423051 | CNV-gain | 1 | CC-NBS-LRR |
| LOC_Os06g33360 | 19434822-19439087 | CNV-gain | 1 | NBS |
| LOC_Os06g43670 | 26279804-26292345 | CNV-gain | 1 | CC-NBS-LRR |
| LOC_Os06g45690 | 27666205-27670507 | CNV-gain | 1 | NBS-LRR |
| LOC_Os06g45820 | 27724565-27728316 | CNV-gain | 1 | NBS |
| LOC_Os06g48520 | 29352930-29356136 | CNV-gain | 1 | NBS-LRR |
| LOC_Os06g49380 | 29917712-29924540 | CNV-loss | 1 | NBS-LRR |
| LOC_Os06g50050 | 30332479-30336110 | CNV-gain | 1 | NBS |
| LOC_Os07g01530 | 333160-334829 | CNV-gain | 1 | NBS |
| LOC_Os07g02530 | 899100-899779 | CNV-gain | 1 | NBS |
| LOC_Os07g02560 | 919573-921478 | CNV-gain | 1 | CC-NBS |
| LOC_Os07g02570 | 925306-926838 | CNV-gain | 2 | NBS |
| LOC_Os07g02590 | 928495-931304 | CNV-gain | 2 | NBS |
| LOC_Os07g02610 | 939612-941748 | CNV-gain | 2 | NBS |
| LOC_Os07g02620 | 945207-946778 | CNV-gain | 2 | CC-NBS |
| LOC_Os07g02630 | 950862-953907 | CNV-loss | 2 | NBS |
| LOC_Os07g02640 | 956244-957651 | CNV-loss | 2 | NBS |
| LOC_Os07g06644 | 3241936-3244694 | CNV-gain | 1 | NBS |
| LOC_Os07g12670 | 7239161-7241105 | CNV-loss | 1 | NBS |
| LOC_Os07g12680 | 7244144-7245017 | CNV-loss | 1 | NBS |
| LOC_Os07g17250 | 10173157-10176827 | CNV-loss | 1 | NBS-LRR |
| LOC_Os07g29810 | 17524037-17528680 | CNV-gain | 1 | NBS-LRR |
| LOC_Os08g01580 | 342215-346733 | CNV-gain | 1 | NBS-LRR |
| LOC_Os08g07330 | 4105398-4109958 | CNV-loss | 1 | NBS-LRR |
| LOC_Os08g07340 | 4111868-4115417 | CNV-loss | 1 | NBS-LRR |

| | | | | |
|---|---|---|---|---|
| LOC_Os08g07774 | 4352807-4361458 | CNV-loss/gain | 2 | NBS-LRR |
| LOC_Os08g07890 | 4440181-4448191 | CNV-loss | 1 | NBS-LRR |
| LOC_Os08g07920 | 4462149-4464256 | CNV-loss | 1 | CC-NBS |
| LOC_Os08g07930 | 4466029-4471773 | CNV-loss | 1 | NBS-LRR |
| LOC_Os08g07940 | 4474830-4485343 | CNV-loss | 1 | NBS-LRR |
| LOC_Os08g07950 | 4487727-4495598 | CNV-gain | 1 | NBS-LRR |
| LOC_Os08g09430 | 5466374-5469304 | CNV-gain | 1 | NBS-LRR |
| LOC_Os08g10430 | 6127418-6131390 | CNV-gain | 1 | NBS-LRR |
| LOC_Os08g24380 | 14714652-14721326 | CNV-gain | 1 | NBS-LRR |
| LOC_Os08g31780 | 19701779-19705239 | CNV-loss | 2 | CC-NBS-LRR |
| LOC_Os08g31800 | 19715508-19719680 | CNV-gain | 1 | NBS-LRR |
| LOC_Os08g32880 | 20389564-20392287 | CNV-gain | 1 | NBS-LRR |
| LOC_Os08g32890 | 20395487-20398504 | CNV-gain | 1 | NBS-LRR |
| LOC_Os08g38970 | 24633408-24637194 | CNV-gain | 1 | NBS |
| LOC_Os08g42670 | 26968542-26978241 | CNV-gain | 1 | CC-NBS-LRR |
| LOC_Os08g42710 | 27016962-27027229 | CNV-loss | 1 | NBS-LRR |
| LOC_Os08g43000 | 27175631-27179263 | CNV-loss | 1 | CC-NBS-LRR |
| LOC_Os08g43010 | 27181459-27185448 | CNV-loss | 1 | NBS-LRR |
| LOC_Os08g43050 | 27205869-27209951 | CNV-gain | 1 | CC-NBS-LRR |
| LOC_Os08g44240 | 27849043-27856246 | CNV-gain | 1 | NBS |
| LOC_Os08g45010 | 28250419-28254009 | CNV-gain | 1 | NBS |
| LOC_Os09g14450 | 8537185-8541654 | CNV-gain | 1 | NBS-LRR |
| LOC_Os09g14490 | 8564278-8572456 | CNV-gain | 1 | NBS-LRR |
| LOC_Os09g20020 | 11989748-11993428 | CNV-gain | 1 | NBS-LRR |
| LOC_Os09g20040 | 12004419-12010546 | CNV-loss | 1 | NBS-LRR |
| LOC_Os09g30220 | 18391869-18397839 | CNV-gain | 1 | NBS-LRR |
| LOC_Os09g34150 | 20157700-20162008 | CNV-gain | 1 | NBS-LRR |
| LOC_Os09g36300 | 20945915-20952936 | CNV-gain | 1 | NBS |
| LOC_Os10g04060 | 1871009-1875535 | CNV-gain | 1 | NBS-LRR |
| LOC_Os10g04674 | 2226402-2237611 | CNV-gain | 1 | NBS-LRR |
| LOC_Os10g07534 | 4026942-4029695 | CNV-gain | 2 | NBS-LRR |
| LOC_Os10g17690 | 8939933-8942517 | CNV-gain | 1 | NBS-LRR |
| LOC_Os10g36270 | 19389025-19396741 | CNV-gain | 1 | NBS-LRR |

| | | | | |
|---|---|---|---|---|
| LOC_Os11g03650 | 1416436-1421232 | CNV-gain | 1 | NBS-LRR |
| LOC_Os11g06300 | 3023705-3025129 | CNV-gain | 2 | NBS |
| LOC_Os11g10120 | 5467512-5469283 | CNV-gain | 1 | NBS |
| LOC_Os11g10770 | 5911937-5919029 | CNV-gain | 1 | NBS |
| LOC_Os11g11550 | 6419845-6424350 | CNV-loss | 2 | NBS-LRR |
| LOC_Os11g11580 | 6440319-6449529 | CNV-loss | 2 | NBS-LRR |
| LOC_Os11g11790 | 6541924-6546026 | CNV-gain | 1 | NBS-LRR |
| LOC_Os11g11810 | 6554514-6561687 | CNV-loss | 1 | NBS-LRR |
| LOC_Os11g11920 | 6606427-6616027 | CNV-loss | 1 | NBS-LRR |
| LOC_Os11g11940 | 6621642-6629013 | CNV-loss | 1 | NBS |
| LOC_Os11g11950 | 6631958-6634662 | CNV-loss/gain | 2 | NBS-LRR |
| LOC_Os11g11960 | 6638262-6640922 | CNV-loss/gain | 2 | NBS-LRR |
| LOC_Os11g13410 | 7342091-7345984 | CNV-loss/gain | 2 | NBS-LRR |
| LOC_Os11g13430 | 7356166-7359604 | CNV-gain | 1 | CC-NBS-LRR |
| LOC_Os11g13440 | 7365098-7367861 | CNV-gain | 1 | CC-NBS-LRR |
| LOC_Os11g29000 | 16781662-16784646 | CNV-loss | 1 | NBS-LRR |
| LOC_Os11g29014 | 16791974-16796861 | CNV-loss | 1 | CC-NBS |
| LOC_Os11g29030 | 16807964-16809227 | CNV-loss | 1 | NBS |
| LOC_Os11g29050 | 16821579-16826538 | CNV-loss | 1 | NBS-LRR |
| LOC_Os11g29110 | 16860235-16864216 | CNV-loss | 1 | NBS-LRR |
| LOC_Os11g29920 | 17385302-17393700 | CNV-gain | 1 | CC-NBS-LRR |
| LOC_Os11g29970 | 17411995-17417481 | CNV-gain | 2 | NBS-LRR |
| LOC_Os11g29980 | 17425545-17429007 | CNV-gain | 2 | NBS |
| LOC_Os11g29990 | 17434258-17438286 | CNV-gain | 2 | NBS-LRR |
| LOC_Os11g30050 | 17479444-17483263 | CNV-gain | 1 | NBS |
| LOC_Os11g30060 | 17484860-17489555 | CNV-gain | 1 | NBS-LRR |
| LOC_Os11g32170 | 18990892-18999281 | CNV-gain | 1 | CC-NBS-LRR |
| LOC_Os11g32210 | 19022758-19029390 | CNV-gain | 1 | CC-NBS-LRR |
| LOC_Os11g34350 | 20128987-20134345 | CNV-gain | 1 | NBS |
| LOC_Os11g36760 | 21705480-21708029 | CNV-loss | 1 | NBS |
| LOC_Os11g37050 | 21872063-21877085 | CNV-loss | 1 | NBS-LRR |
| LOC_Os11g37260 | 22003368-22010520 | CNV-gain | 1 | NBS |
| LOC_Os11g38580 | 22862447-22867268 | CNV-loss | 1 | CC-NBS-LRR |

| | | | | |
|---|---|---|---|---|
| LOC_Os11g39020 | 23230112-23234077 | CNV-gain | 1 | CC-NBS |
| LOC_Os11g39250 | 23368577-23373607 | CNV-gain | 1 | NBS |
| LOC_Os11g39330 | 23415625-23418638 | CNV-gain | 1 | NBS-LRR |
| LOC_Os11g40780 | 24387280-24390444 | CNV-loss | 1 | CC-NBS-LRR |
| LOC_Os11g41540 | 24911501-24917050 | CNV-gain | 1 | NBS |
| LOC_Os11g42040 | 25289129-25293954 | CNV-gain | 1 | CC-NBS-LRR |
| LOC_Os11g42070 | 25321310-25325479 | CNV-loss | 1 | NBS-LRR |
| LOC_Os11g42090 | 25338882-25345347 | CNV-loss | 1 | NBS-LRR |
| LOC_Os11g42100 | 25358357-25360219 | CNV-loss | 1 | NBS-LRR |
| LOC_Os11g43250 | 26088067-26091283 | CNV-loss | 1 | CC-NBS-LRR |
| LOC_Os11g43320 | 26139957-26143460 | CNV-loss | 1 | NBS-LRR |
| LOC_Os11g44960 | 27221855-27226087 | CNV-loss/gain | 2 | CC-NBS-LRR |
| LOC_Os11g44970 | 27234199-27237260 | CNV-gain | 1 | CC-NBS-LRR |
| LOC_Os11g44990 | 27242070-27245161 | CNV-loss | 2 | CC-NBS-LRR |
| LOC_Os11g45050 | 27261093-27264194 | CNV-loss | 1 | CC-NBS-LRR |
| LOC_Os11g45090 | 27282232-27285045 | CNV-gain | 1 | CC-NBS-LRR |
| LOC_Os11g45130 | 27318942-27324163 | CNV-gain | 1 | NBS-LRR |
| LOC_Os11g45160 | 27335214-27337380 | CNV-gain | 1 | CC-NBS |
| LOC_Os11g45180 | 27342028-27345123 | CNV-gain | 1 | CC-NBS-LRR |
| LOC_Os11g45330 | 27427776-27430560 | CNV-loss | 1 | NBS-LRR |
| LOC_Os12g09240 | 4836884-4841595 | CNV-loss | 1 | CC-NBS-LRR |
| LOC_Os12g13550 | 7602046-7606922 | CNV-gain | 1 | NBS-LRR |
| LOC_Os12g30590 | 18369705-18373748 | CNV-gain | 1 | NBS |
| LOC_Os12g30720 | 18446075-18450017 | CNV-gain | 1 | CC-NBS-LRR |
| LOC_Os12g30760 | 18469213-18474449 | CNV-loss/gain | 2 | CC-NBS-LRR |
| LOC_Os12g31620 | 19043758-19048583 | CNV-loss | 2 | NBS-LRR |
| LOC_Os12g32590 | 19663107-19667757 | CNV-gain | 1 | NBS |
| LOC_Os12g32660 | 19703788-19710182 | CNV-gain | 1 | NBS-LRR |
| LOC_Os12g32680 | 19730411-19734112 | CNV-gain | 1 | NBS |
| LOC_Os12g32710 | 19754146-19757913 | CNV-loss | 1 | NBS-LRR |
| LOC_Os12g33160 | 20061564-20066930 | CNV-gain | 1 | CC-NBS-LRR |
| LOC_Os12g36690 | 22477070-22481287 | CNV-loss | 2 | CC-NBS |
| LOC_Os12g36720 | 22487337-22492402 | CNV-loss | 2 | CC-NBS-LRR |

| | | | | |
|---|---|---|---|---|
| LOC_Os12g36730 | 22497534-22502497 | CNV-loss | 1 | CC-NBS-LRR |
| LOC_Os12g40880 | 25305806-25309780 | CNV-gain | 1 | NBS |

**Table S28. Summary of PAV calling between *O. sativa* and *O. rufipogon*, *O. sativa* and *O. nivara*, respectively.** PAV types are represented in a customized format, of which RS10 indicates that a PAV is present RUF but absent in SAT, NS10 indicates that a PAV is present NIV but absent in SAT, RS01 indicates that a PAV is absent RUF but present in SAT, and NS01 indicates a PAV is absent RUF but present in SAT.

| **PAV Type** | **Number of PAVs** | | | | | | |
|---|---|---|---|---|---|---|---|
| | 100-500 | 501-1000 | 1001-2000 | 2001-5000 | 5001-20000 | >20000 | **Total** |
| **RS10** | 11,255 | 5,310 | 4,362 | 3,236 | 1,198 | 26 | **25,387** |
| **RS01** | 5,773 | 1,845 | 1,148 | 1,143 | 599 | 11 | **10,519** |
| **NS10** | 6,448 | 2,888 | 2,431 | 1,962 | 702 | 8 | **14,439** |
| **NS01** | 19,735 | 7,477 | 4,192 | 2,670 | 1,083 | 24 | **35,181** |

**Table S29. Summary of PAV calling between *O. sativa* and *O. rufipogon*, *O. sativa* and *O. nivara*, respectively.**

| PAV Type | Length of PAV regions (bp) | Gene number in PAV regions | Gene number with known functions |
|---|---|---|---|
| **RS10** | 16,232,326 | 4,621 | 3,018 |
| **RS01** | 9,503,214 | 3,241 | 1,123 |
| **NS10** | 9,161,166 | 3,076 | 1,701 |
| **NS01** | 23,220,789 | 14,425 | 6,906 |

**Note:** R: *O. rufipogon*; S: ; N *O. sativa*: *O. nivara*; 1: presence; 0: absence.

**Table S30. PFAM functional enrichment analysis of genes associated with PAVs.** The top 10 Pfam IDs are given for each PAV type. Significance levels of functional enrichment are shown for *P*-value and FDR.

| PAV | PFAM | Description | #Gene | *P*-value | FDR |
|---|---|---|---|---|---|
| **RS10** | PF00931 | NB-ARC domain | 187 | 4.89E-43 | 4.34E-41 |
| **RS10** | PF00069 | Protein kinase domain | 154 | 1.80E-02 | 2.32E-01 |
| **RS10** | PF07714 | Protein tyrosine kinase | 81 | 5.08E-04 | 3.00E-02 |
| **RS10** | PF00067 | Cytochrome P450 | 57 | 7.51E-02 | 2.32E-01 |
| **RS10** | PF03140 | Plant protein of unknown function | 46 | 1.22E-18 | 1.06E-16 |
| **RS10** | PF13855 | Leucine rich repeat | 43 | 6.51E-01 | 7.27E-01 |
| **RS10** | PF13947 | Wall-associated receptor kinase galacturonan-binding | 40 | 2.33E-08 | 1.99E-06 |
| **RS10** | PF00560 | Leucine Rich Repeat | 37 | 9.62E-01 | 9.78E-01 |
| **RS10** | PF13968 | Domain of unknown function (DUF4220) | 36 | 3.47E-08 | 2.90E-06 |
| **RS10** | PF00646 | F-box domain | 36 | 6.07E-01 | 6.90E-01 |
| **RS01** | PF00931 | NB-ARC domain | 63 | 5.46E-14 | 8.19E-12 |
| **RS01** | PF00069 | Protein kinase domain | 46 | 5.41E-01 | 5.67E-01 |
| **RS01** | PF00067 | Cytochrome P450 | 27 | 5.50E-03 | 1.22E-01 |
| **RS01** | PF07714 | Protein tyrosine kinase | 24 | 4.19E-01 | 4.59E-01 |
| **RS01** | PF00847 | AP2 domain | 14 | 7.05E-03 | 1.22E-01 |
| **RS01** | PF00201 | UDP-glucoronosyl and UDP-glucosyl transferase | 13 | 8.09E-02 | 1.63E-01 |
| **RS01** | PF13968 | Domain of unknown function (DUF4220) | 12 | 1.58E-03 | 1.22E-01 |
| **RS01** | PF13855 | Leucine rich repeat | 12 | 9.27E-01 | 9.31E-01 |
| **RS01** | PF13041 | PPR repeat family | 12 | 9.61E-01 | 9.64E-01 |
| **RS01** | PF00249 | Myb-like DNA-binding domain | 11 | 3.79E-01 | 4.27E-01 |
| **NS10** | PF00931 | NB-ARC domain | 122 | 2.20E-33 | 3.18E-31 |
| **NS10** | PF00069 | Protein kinase domain | 88 | 1.18E-01 | 2.59E-01 |
| **NS10** | PF00067 | Cytochrome P450 | 51 | 9.96E-08 | 1.29E-05 |
| **NS10** | PF00646 | F-box domain | 32 | 1.88E-03 | 1.47E-01 |
| **NS10** | PF07714 | Protein tyrosine kinase | 30 | 6.39E-01 | 6.85E-01 |
| **NS10** | PF13968 | Domain of unknown function (DUF4220) | 28 | 1.66E-08 | 2.22E-06 |
| **NS10** | PF13855 | Leucine rich repeat | 24 | 7.19E-01 | 7.56E-01 |
| **NS10** | PF03140 | Plant protein of unknown function | 23 | 9.87E-09 | 1.37E-06 |
| **NS10** | PF00560 | Leucine Rich Repeat | 23 | 8.31E-01 | 8.54E-01 |
| **NS10** | PF13947 | Wall-associated receptor kinase galacturonan-binding | 22 | 5.19E-06 | 6.29E-04 |
| **NS01** | PF00069 | Protein kinase domain | 200 | 1.00E+00 | 1.00E+00 |
| **NS01** | PF00067 | Cytochrome P450 | 147 | 1.58E-07 | 2.13E-06 |
| **NS01** | PF13639 | Ring finger domain | 121 | 1.85E-08 | 2.53E-07 |
| **NS01** | PF00931 | NB-ARC domain | 112 | 9.97E-01 | 1.00E+00 |
| **NS01** | PF07714 | Protein tyrosine kinase | 103 | 1.00E+00 | 1.00E+00 |
| **NS01** | PF00847 | AP2 domain | 99 | 9.14E-20 | 1.27E-18 |
| **NS01** | PF00646 | F-box domain | 94 | 7.25E-01 | 8.83E-01 |

| | | | | | |
|---|---|---|---|---|---|
| **NS01** | PF00201 | UDP-glucoronosyl and UDP-glucosyl transferase | 91 | 7.72E-08 | 1.05E-06 |
| **NS01** | PF00249 | Myb-like DNA-binding domain | 78 | 3.01E-02 | 2.80E-01 |
| **NS01** | PF02458 | Transferase family | 73 | 7.38E-11 | 1.02E-09 |

**Table S31. Gene Ontology (GO) functional enrichment analysis of genes associated with PAVs.** The top 10 GO IDs are given below for each PAV type. Significance levels of functional enrichment are shown for *P*-value and FDR.

| PAV | GO | Description | #Gene | *P*-value | FDR |
|---|---|---|---|---|---|
| **RS10** | GO:0004672 | protein kinase activity | 279 | 5.78E-20 | 3.93E-18 |
| **RS10** | GO:0005515 | protein binding | 224 | 9.77E-01 | 9.95E-01 |
| **RS10** | GO:0043531 | ADP binding | 203 | 4.54E-54 | 3.43E-52 |
| **RS10** | GO:0008152 | metabolic process | 71 | 2.22E-01 | 4.33E-01 |
| **RS10** | GO:0003677 | DNA binding | 66 | 9.16E-01 | 9.54E-01 |
| **RS10** | GO:0005506 | iron ion binding | 59 | 3.15E-01 | 4.99E-01 |
| **RS10** | GO:0003676 | nucleic acid binding | 51 | 8.82E-01 | 9.27E-01 |
| **RS10** | GO:0016021 | integral component of membrane | 51 | 9.21E-01 | 9.56E-01 |
| **RS10** | GO:0003824 | catalytic activity | 50 | 8.81E-01 | 9.27E-01 |
| **RS10** | GO:0016491 | oxidoreductase activity | 45 | 1.12E-01 | 3.47E-01 |
| **RS01** | GO:0005515 | protein binding | 22 | 3.62E-01 | 4.27E-01 |
| **RS01** | GO:0004672 | protein kinase activity | 103 | 2.10E-05 | 2.68E-03 |
| **RS01** | GO:0043531 | ADP binding | 84 | 9.92E-15 | 1.58E-12 |
| **RS01** | GO:0008152 | metabolic process | 63 | 4.82E-02 | 1.39E-01 |
| **RS01** | GO:0005506 | iron ion binding | 34 | 6.06E-03 | 1.37E-01 |
| **RS01** | GO:0003700 | sequence-specific DNA binding transcription factor activity | 29 | 3.29E-02 | 1.37E-01 |
| **RS01** | GO:0003677 | DNA binding | 28 | 9.09E-01 | 9.18E-01 |
| **RS01** | GO:0003824 | catalytic activity | 21 | 6.72E-01 | 7.10E-01 |
| **RS01** | GO:0004553 | hydrolase activity, hydrolyzing O-glycosyl compounds | 14 | 2.65E-01 | 3.41E-01 |
| **RS01** | GO:0016491 | oxidoreductase activity | 13 | 4.10E-01 | 4.61E-01 |
| **NS10** | GO:0004672 | protein kinase activity | 160 | 2.00E-12 | 1.77E-10 |
| **NS10** | GO:0005515 | protein binding | 147 | 1.98E-01 | 3.52E-01 |
| **NS10** | GO:0043531 | ADP binding | 138 | 2.62E-46 | 2.70E-44 |
| **NS10** | GO:0005506 | iron ion binding | 51 | 1.50E-05 | 9.25E-04 |
| **NS10** | GO:0008152 | metabolic process | 44 | 9.16E-03 | 2.37E-01 |
| **NS10** | GO:0003677 | DNA binding | 36 | 9.78E-01 | 9.90E-01 |
| **NS10** | GO:0003824 | catalytic activity | 29 | 7.91E-01 | 8.34E-01 |
| **NS10** | GO:0016491 | oxidoreductase activity | 28 | 9.77E-02 | 2.89E-01 |
| **NS10** | GO:0003676 | nucleic acid binding | 25 | 9.80E-01 | 9.91E-01 |
| **NS10** | GO:0030247 | polysaccharide binding | 22 | 4.80E-11 | 3.70E-09 |
| **NS01** | GO:0005515 | protein binding | 673 | 3.25E-03 | 2.88E-02 |
| **NS01** | GO:0004672 | protein kinase activity | 346 | 2.26E-01 | 5.51E-01 |
| **NS01** | GO:0008152 | metabolic process | 243 | 2.29E-14 | 2.44E-13 |
| **NS01** | GO:0003700 | sequence-specific DNA binding transcription factor activity | 215 | 5.78E-21 | 6.25E-20 |
| **NS01** | GO:0003677 | DNA binding | 169 | 6.57E-01 | 8.22E-01 |

| | | | | | |
|---|---|---|---|---|---|
| **NS01** | GO:0005506 | iron ion binding | 152 | 1.06E-05 | 1.09E-04 |
| **NS01** | GO:0043531 | ADP binding | 114 | 9.88E-01 | 1.00E+00 |
| **NS01** | GO:0005524 | ATP binding | 101 | 2.86E-01 | 5.51E-01 |
| **NS01** | GO:0003824 | catalytic activity | 98 | 6.14E-01 | 8.19E-01 |
| **NS01** | GO:0016491 | oxidoreductase activity | 96 | 1.17E-02 | 9.41E-02 |

**Table S32. Clustering of gene families among the three *Oryza* species, *O. sativa*, *O. rufipogon* and *O. nivara*.**

| Species | Gene number | Genes in families | Unclustered genes | Family number | Unique families | Average gene number per family |
|---|---|---|---|---|---|---|
| **RUF** | 48,445 | 36,902 | 11,543 | 25,391 | 1,007 | 1.45 |
| **NIV** | 38,881 | 29,911 | 8,970 | 24,095 | 437 | 1.24 |
| **SAT** | 39,043 | 33,425 | 5,618 | 26,043 | 239 | 1.28 |

**Table S35. Summary of lineage-specific gene families identified among *O. sativa*, *O. rufipogon*, *O. nivara* and *O. meridionalis*.**

| Species | Status | PFAM_ID | Description | #Gene | *P*-value | FDR |
|---|---|---|---|---|---|---|
| **SAT** | Expansion | PF00428 | 60s Acidic ribosomal protein | 3 | 3.04E-05 | 3.95E-03 |
| **SAT** | Expansion | PF13087 | AAA domain | 2 | 2.02E-03 | 4.61E-02 |
| **SAT** | Expansion | PF01536 | Adenosylmethionine decarboxylase | 5 | 1.41E-09 | 5.00E-07 |
| **SAT** | Expansion | PF01263 | Aldose 1-epimerase | 3 | 4.33E-05 | 5.45E-03 |
| **NIV** | Expansion | PF12695 | Alpha/beta hydrolase family | 8 | 1.01E-26 | 3.79E-23 |
| **SAT** | Expansion | PF00306 | ATP synthase alpha/beta chain, C terminal domain | 4 | 5.75E-06 | 9.35E-04 |
| **SAT** | Expansion | PF00006 | ATP synthase alpha/beta family, nucleotide-binding domain | 3 | 1.74E-04 | 2.00E-02 |
| **SAT** | Expansion | PF00497 | Bacterial extracellular solute-binding proteins, family 3 | 2 | 4.53E-03 | 4.61E-02 |
| **NIV** | Contraction | PF07645 | Calcium-binding EGF domain | 5 | 1.64E-04 | 4.62E-02 |
| **RUF** | Expansion | PF07645 | Calcium-binding EGF domain | 16 | 1.51E-09 | 3.07E-07 |
| **NIV** | Contraction | PF12819 | Carbohydrate-binding protein of the ER | 5 | 3.78E-04 | 4.62E-02 |
| **SAT** | Expansion | PF12819 | Carbohydrate-binding protein of the ER | 14 | 6.46E-14 | 8.40E-11 |
| **RUF** | Expansion | PF02386 | Cation transport protein | 8 | 7.48E-08 | 1.23E-05 |
| **SAT** | Expansion | PF12348 | CLASP N terminal | 3 | 4.52E-06 | 7.67E-04 |
| **RUF** | Expansion | PF01656 | CobQ/CobB/MinD/ParA nucleotide binding domain | 3 | 9.40E-05 | 9.54E-03 |
| **RUF** | Expansion | PF00125 | Core histone H2A/H2B/H3/H4 | 15 | 5.53E-07 | 7.14E-05 |
| **RUF** | Expansion | PF02672 | CP12 domain | 4 | 1.71E-05 | 1.92E-03 |
| **SAT** | Expansion | PF13016 | Cys-rich Gliadin N-terminal | 5 | 1.05E-06 | 2.04E-04 |
| **SAT** | Expansion | PF00032 | Cytochrome b(C-terminal)/b6/petD | 3 | 6.80E-08 | 1.77E-05 |
| **NIV** | Contraction | PF00067 | Cytochrome P450 | 13 | 3.65E-04 | 4.62E-02 |
| **RUF** | Expansion | PF00067 | Cytochrome P450 | 78 | 1.00E-14 | 3.88E-12 |
| **SAT** | Expansion | PF00067 | Cytochrome P450 | 13 | 2.06E-03 | 4.61E-02 |
| **NIV** | Contraction | PF01453 | D-mannose binding lectin | 11 | 1.03E-07 | 4.31E-05 |
| **RUF** | Expansion | PF01453 | D-mannose binding lectin | 27 | 9.31E-08 | 1.47E-05 |
| **NIV** | Contraction | PF08224 | Domain of unknown function (DUF1719) | 3 | 1.63E-06 | 6.13E-04 |
| **RUF** | Expansion | PF09322 | Domain of unknown function (DUF1979) | 36 | 1.39E-33 | 9.87E-31 |
| **RUF** | Expansion | PF13968 | Domain of unknown function (DUF4220) | 23 | 8.77E-06 | 1.01E-03 |
| **RUF** | Expansion | PF14111 | Domain of unknown function (DUF4283) | 10 | 3.57E-05 | 3.80E-03 |
| **SAT** | Expansion | PF14291 | Domain of unknown function (DUF4371) | 9 | 4.71E-12 | 3.68E-09 |
| **RUF** | Expansion | PF14365 | Domain of unknown | 10 | 4.13E-07 | 5.86E-05 |

| | | | | | | |
|---|---|---|---|---|---|---|
| | | | function (DUF4409) | | | |
| **SAT** | Expansion | PF05754 | Domain of unknown function (DUF834) | 5 | 3.81E-04 | 3.72E-02 |
| **SAT** | Expansion | PF01266 | FAD dependent oxidoreductase | 3 | 8.04E-06 | 1.21E-03 |
| **RUF** | Expansion | PF01151 | GNS1/SUR4 family | 7 | 1.89E-08 | 3.51E-06 |
| **RUF** | Expansion | PF03514 | GRAS domain family | 20 | 1.00E-10 | 2.37E-08 |
| **RUF** | Expansion | PF01493 | GXGXG motif | 2 | 5.13E-04 | 4.56E-02 |
| **SAT** | Expansion | PF05699 | hAT family C-terminal dimerisation region | 3 | 1.06E-04 | 1.25E-02 |
| **RUF** | Expansion | PF14214 | Helitron helicase-like domain at N-terminus | 27 | 1.59E-35 | 1.35E-32 |
| **SAT** | Contraction | PF14214 | Helitron helicase-like domain at N-terminus | 2 | 1.27E-10 | 2.48E-07 |
| **SAT** | Expansion | PF00742 | Homoserine dehydrogenase | 2 | 1.67E-05 | 2.33E-03 |
| **RUF** | Expansion | PF02301 | HORMA domain | 7 | 1.89E-08 | 3.51E-06 |
| **SAT** | Expansion | PF01419 | Jacalin-like lectin domain | 2 | 1.03E-02 | 4.61E-02 |
| **NIV** | Contraction | PF00139 | Legume lectin domain | 20 | 3.19E-21 | 5.98E-18 |
| **RUF** | Expansion | PF00139 | Legume lectin domain | 26 | 2.07E-11 | 5.89E-09 |
| **NIV** | Contraction | PF13855 | Leucine rich repeat | 29 | 5.31E-14 | 3.32E-11 |
| **NIV** | Contraction | PF00560 | Leucine Rich Repeat | 15 | 2.10E-04 | 4.62E-02 |
| **NIV** | Contraction | PF13504 | Leucine rich repeat | 3 | 4.10E-03 | 4.62E-02 |
| **RUF** | Expansion | PF00560 | Leucine Rich Repeat | 62 | 1.07E-07 | 1.63E-05 |
| **RUF** | Expansion | PF13855 | Leucine rich repeat | 56 | 1.49E-06 | 1.81E-04 |
| **NIV** | Contraction | PF08263 | Leucine rich repeat N-terminal domain | 15 | 1.82E-06 | 6.22E-04 |
| **NIV** | Contraction | PF12799 | Leucine Rich repeats (2 copies) | 10 | 1.98E-04 | 4.62E-02 |
| **RUF** | Expansion | PF12799 | Leucine Rich repeats (2 copies) | 35 | 5.47E-07 | 7.14E-05 |
| **NIV** | Contraction | PF00060 | Ligand-gated ion channel | 2 | 1.05E-03 | 4.62E-02 |
| **SAT** | Expansion | PF00060 | Ligand-gated ion channel | 3 | 2.17E-04 | 2.42E-02 |
| **SAT** | Expansion | PF08311 | Mad3/BUB1 homology region 1 | 2 | 4.23E-06 | 7.50E-04 |
| **SAT** | Expansion | PF07690 | Major Facilitator Superfamily | 10 | 2.96E-10 | 1.44E-07 |
| **SAT** | Expansion | PF01554 | MatE | 6 | 2.90E-05 | 3.90E-03 |
| **SAT** | Expansion | PF03999 | Microtubule associated protein (MAP65/ASE1 family) | 2 | 4.66E-04 | 4.44E-02 |
| **RUF** | Expansion | PF03108 | MuDR family transposase | 40 | 1.27E-19 | 5.41E-17 |
| **RUF** | Expansion | PF10551 | MULE transposase domain | 24 | 3.32E-11 | 8.83E-09 |
| **SAT** | Expansion | PF12776 | Myb/SANT-like DNA-binding domain | 6 | 4.12E-07 | 8.92E-05 |
| **SAT** | Expansion | PF01370 | NAD dependent epimerase/dehydratase family | 7 | 1.12E-05 | 1.62E-03 |
| **NIV** | Contraction | PF00931 | NB-ARC domain | 36 | 9.78E-16 | 9.18E-13 |
| **RUF** | Expansion | PF00931 | NB-ARC domain | 218 | 6.27E-100 | 6.68E-97 |
| **SAT** | Expansion | PF00931 | NB-ARC domain | 85 | 1.23E-64 | 2.39E-61 |
| **RUF** | Expansion | PF14303 | No apical meristem-associated C-terminal domain | 8 | 3.90E-04 | 3.66E-02 |

| | | | | | | |
|---|---|---|---|---|---|---|
| **SAT** | Expansion | PF14303 | No apical meristem-associated C-terminal domain | 3 | 3.27E-04 | 3.27E-02 |
| **RUF** | Expansion | PF03169 | OPT oligopeptide transporter protein | 26 | 3.36E-24 | 2.04E-21 |
| **NIV** | Contraction | PF08276 | PAN-like domain | 15 | 2.99E-15 | 2.25E-12 |
| **RUF** | Expansion | PF08276 | PAN-like domain | 25 | 1.19E-09 | 2.66E-07 |
| **SAT** | Expansion | PF00141 | Peroxidase | 13 | 2.94E-07 | 7.17E-05 |
| **SAT** | Expansion | PF00124 | Photosynthetic reaction centre protein | 5 | 4.76E-10 | 2.06E-07 |
| **SAT** | Expansion | PF00223 | Photosystem I psaA/psaB protein | 8 | 0.00E+00 | 0.00E+00 |
| **RUF** | Expansion | PF05970 | PIF1-like helicase | 19 | 8.70E-20 | 4.12E-17 |
| **RUF** | Expansion | PF10536 | Plant mobile domain | 44 | 2.64E-21 | 1.40E-18 |
| **SAT** | Contraction | PF10536 | Plant mobile domain | 2 | 4.13E-07 | 4.03E-04 |
| **SAT** | Expansion | PF10536 | Plant mobile domain | 4 | 7.89E-05 | 9.62E-03 |
| **SAT** | Expansion | PF00321 | Plant thionin | 6 | 2.06E-10 | 1.15E-07 |
| **RUF** | Expansion | PF03004 | Plant transposase (Ptta/En/Spm family) | 11 | 4.52E-07 | 6.21E-05 |
| **SAT** | Expansion | PF04928 | Poly(A) polymerase central domain | 2 | 8.34E-04 | 4.61E-02 |
| **RUF** | Expansion | PF11831 | pre-mRNA splicing factor component | 2 | 0.00E+00 | 0.00E+00 |
| **NIV** | Contraction | PF00069 | Protein kinase domain | 70 | 2.92E-27 | 1.10E-23 |
| **RUF** | Expansion | PF00069 | Protein kinase domain | 133 | 4.76E-08 | 8.12E-06 |
| **SAT** | Expansion | PF00069 | Protein kinase domain | 30 | 4.87E-04 | 4.52E-02 |
| **SAT** | Expansion | PF07762 | Protein of unknown function (DUF1618) | 7 | 2.58E-04 | 2.69E-02 |
| **RUF** | Expansion | PF11145 | Protein of unknown function (DUF2921) | 8 | 1.39E-09 | 2.96E-07 |
| **SAT** | Expansion | PF12425 | Protein of unknown function (DUF3673) | 5 | 4.70E-08 | 1.53E-05 |
| **RUF** | Expansion | PF12527 | Protein of unknown function (DUF3727) | 3 | 3.27E-05 | 3.57E-03 |
| **SAT** | Contraction | PF04578 | Protein of unknown function, DUF594 | 2 | 1.99E-05 | 1.61E-03 |
| **NIV** | Contraction | PF07714 | Protein tyrosine kinase | 38 | 2.68E-19 | 3.35E-16 |
| **RUF** | Expansion | PF04195 | Putative gypsy type transposon | 25 | 1.10E-12 | 3.89E-10 |
| **RUF** | Expansion | PF01170 | Putative RNA methylase family UPF0020 | 3 | 6.81E-06 | 8.06E-04 |
| **NIV** | Contraction | PF01094 | Receptor family ligand binding region | 2 | 1.05E-03 | 4.62E-02 |
| **SAT** | Expansion | PF00346 | Respiratory-chain NADH dehydrogenase, 49 Kd subunit | 3 | 6.80E-08 | 1.77E-05 |
| **RUF** | Expansion | PF00078 | Reverse transcriptase (RNA-dependent DNA polymerase) | 11 | 5.20E-05 | 5.40E-03 |
| **SAT** | Expansion | PF07727 | Reverse transcriptase (RNA-dependent DNA polymerase) | 3 | 6.80E-08 | 1.77E-05 |
| **NIV** | Expansion | PF00832 | Ribosomal L39 protein | 2 | 2.42E-10 | 4.54E-07 |
| **SAT** | Expansion | PF00252 | Ribosomal protein | 5 | 1.41E-09 | 5.00E-07 |

| | | | | | | |
|---|---|---|---|---|---|---|
| | | | L16p/L10e | | | |
| **SAT** | Expansion | PF00237 | Ribosomal protein L22p/L17e | 3 | 2.29E-06 | 4.26E-04 |
| **RUF** | Expansion | PF01246 | Ribosomal protein L24e | 3 | 4.04E-04 | 3.66E-02 |
| **RUF** | Expansion | PF01247 | Ribosomal protein L35Ae | 3 | 2.10E-04 | 2.04E-02 |
| **SAT** | Expansion | PF00203 | Ribosomal protein S19 | 6 | 2.20E-12 | 2.15E-09 |
| **SAT** | Expansion | PF00318 | Ribosomal protein S2 | 3 | 8.04E-06 | 1.21E-03 |
| **SAT** | Expansion | PF00410 | Ribosomal protein S8 | 3 | 9.94E-07 | 2.04E-04 |
| **RUF** | Expansion | PF00101 | Ribulose bisphosphate carboxylase, small chain | 3 | 4.04E-04 | 3.66E-02 |
| **RUF** | Expansion | PF09133 | SANTA (SANT Associated) | 2 | 0.00E+00 | 0.00E+00 |
| **NIV** | Contraction | PF00954 | S-locus glycoprotein family | 10 | 2.85E-08 | 1.34E-05 |
| **RUF** | Expansion | PF00954 | S-locus glycoprotein family | 24 | 3.30E-08 | 5.86E-06 |
| **RUF** | Expansion | PF00862 | Sucrose synthase | 9 | 1.21E-07 | 1.78E-05 |
| **RUF** | Expansion | PF00083 | Sugar (and other) transporter | 34 | 8.09E-12 | 2.46E-09 |
| **SAT** | Expansion | PF01103 | Surface antigen | 2 | 2.23E-04 | 2.42E-02 |
| **SAT** | Expansion | PF07719 | Tetratricopeptide repeat | 4 | 7.05E-04 | 4.61E-02 |
| **SAT** | Expansion | PF00290 | Tryptophan synthase alpha chain | 3 | 3.36E-07 | 7.71E-05 |
| **NIV** | Contraction | PF00201 | UDP-glucoronosyl and UDP-glucosyl transferase | 8 | 6.53E-05 | 2.04E-02 |
| **RUF** | Expansion | PF00201 | UDP-glucoronosyl and UDP-glucosyl transferase | 25 | 1.79E-04 | 1.78E-02 |
| **RUF** | Expansion | PF02902 | Ulp1 protease family, C-terminal catalytic domain | 22 | 4.74E-11 | 1.19E-08 |
| **SAT** | Expansion | PF02902 | Ulp1 protease family, C-terminal catalytic domain | 12 | 5.03E-11 | 3.27E-08 |
| **NIV** | Contraction | PF13947 | Wall-associated receptor kinase galacturonan-binding | 15 | 8.09E-12 | 4.34E-09 |
| **RUF** | Expansion | PF13947 | Wall-associated receptor kinase galacturonan-binding | 27 | 1.36E-06 | 1.70E-04 |
| **SAT** | Expansion | PF13920 | Zinc finger, C3HC4 type (RING finger) | 3 | 1.54E-02 | 4.61E-02 |
| **RUF** | Expansion | PF13966 | zinc-binding in reverse transcriptase | 21 | 5.24E-12 | 1.72E-09 |

**Table S38. Number of putative NBS-LRR genes identified in the three *Oryza* species, *O. sativa*, *O. rufipogon* and *O. nivara*.**

| | **RUF** | **SAT** | **NIV** |
|---|---|---|---|
| CC-NBS | 76 | 56 | 64 |
| CC-NBS-LRR | 166 | 252 | 133 |
| NBS-LRR | 215 | 227 | 178 |
| TIR-NBS | 1 | 1 | 1 |
| NBS | 118 | 95 | 113 |
| Total | **576** | **631** | **489** |

**Table S39. Summary of branch-specific ω, *dN*, and *dS* values along *O. rufipogon*, *O. sativa*, *O. nivara* and *O. meridionalis* lineages estimated by using PAML.**

| Species/branch[a] | ω(*dN*/*dS*) | *dN* | *dS* |
|---|---|---|---|
| **1# RUF** | 0.5382 | 0.005491 | 0.010202 |
| **2# SAT** | 0.5352 | 0.002995 | 0.005596 |
| **3# NIV** | 0.6598 | 0.009203 | 0.013947 |
| **4# MER** | 0.3618 | 0.014485 | 0.040033 |
| **5# Ancestral of RUF&SAT** | 0.6333 | 0.001826 | 0.002883 |

**Table S40. Summary of the numbers of PSGs detected along *O. rufipogon*, *O. sativa* and *O. nivara* lineages estimated by using PAML.**

| **Branch** | **All PSGs** | | **Lineage-specific PSGs** [c] | |
|---|---|---|---|---|
| | **P < 0.05** | **FDR < 0.05** | **P < 0.05** | **FDR < 0.05** |
| [a] **All branches** | 1,905 | 1,799 | - | - |
| **SAT** | 273 | 247 | 103 | 90 |
| **NIV** | 1,017 | 996 | 487 | 476 |
| **RUF** | 440 | 416 | 211 | 199 |
| [b] **Ancestral of RUF&SAT** | 250 | 235 | 53 | 50 |
| **Total (non-redundancy)** | 2,148 | 2,053 | 854 | 815 |

[a] The genes under selection along any branch of the phylogenetic tree based on the site model;

[b] The branch leading to Asian cultivated rice and *O. rufipogon*;

[c] These genes show significant evidence of positive selection only along one branch.

**Table S47. Functional enrichment analysis of the detected PSGs along *O. rufipogon*, *O. sativa* and *O. nivara* lineages.**

| Type | GO | term | Gene Number | | *P*-values |
|---|---|---|---|---|---|
| | | | PSGs | All | |
| **(A) All non-redundant PSGs** | | | | | |
| MF | GO:0005102 | receptor binding | 3 | 3 | 0 |
| BP | GO:0007610 | behavior | 1 | 1 | 0 |
| BP | GO:0009835 | ripening | 1 | 1 | 0 |
| BP | GO:0009908 | flower development | 65 | 92 | 0.001077 |
| MF | GO:0003700 | sequence-specific DNA binding transcription factor activity | 214 | 346 | 0.007081 |
| BP | GO:0040029 | regulation of gene expression, epigenetic | 10 | 12 | 0.009491 |
| BP | GO:0009607 | response to biotic stimulus | 51 | 79 | 0.043621 |
| BP | GO:0007275 | multicellular organismal development | 144 | 237 | 0.04818 |
| **(B) Clade of RUF&SAT PSGs** | | | | | |
| BP | GO:0007610 | Behavior | 1 | 1 | 0.000531 |
| BP | GO:0040029 | regulation of gene expression, epigenetic | 6 | 12 | 0.001042 |
| CC | GO:0005635 | nuclear envelope | 7 | 18 | 0.002369 |
| BP | GO:0009607 | response to biotic stimulus | 23 | 79 | 0.002604 |
| MF | GO:0019825 | oxygen binding | 5 | 12 | 0.002723 |
| BP | GO:0006412 | Translation | 14 | 45 | 0.003023 |
| BP | GO:0008219 | cell death | 8 | 23 | 0.004745 |
| BP | GO:0009838 | Abscission | 3 | 7 | 0.008073 |
| BP | GO:0030154 | cell differentiation | 19 | 68 | 0.008474 |
| CC | GO:0005739 | Mitochondrion | 39 | 156 | 0.008827 |
| MF | GO:0003824 | catalytic activity | 106 | 466 | 0.012499 |
| BP | GO:0009908 | flower development | 23 | 92 | 0.01648 |
| BP | GO:0000003 | reproduction | 38 | 166 | 0.018238 |
| BP | GO:0009856 | pollination | 2 | 11 | 0.025138 |
| BP | GO:0009991 | response to extracellular stimulus | 6 | 33 | 0.027415 |
| BP | GO:0009791 | post-embryonic development | 35 | 179 | 0.039207 |
| **(C) RUF lineage** | | | | | |
| BP | GO:0007610 | behavior | 1 | 1 | 0 |
| CC | GO:0005623 | cell | 36 | 219 | 0.001228 |
| MF | GO:0019825 | oxygen binding | 4 | 12 | 0.004767 |
| BP | GO:0009719 | response to endogenous stimulus | 23 | 149 | 0.015609 |
| MF | GO:0008135 | translation factor activity, nucleic acid binding | 3 | 12 | 0.027713 |
| MF | GO:0005102 | receptor binding | 1 | 3 | 0.029382 |
| MF | GO:0003824 | catalytic activity | 59 | 466 | 0.030317 |
| BP | GO:0006412 | Translation | 8 | 45 | 0.036003 |
| MF | GO:0004872 | receptor activity | 2 | 8 | 0.040576 |
| BP | GO:0030154 | cell differentiation | 11 | 68 | 0.04137 |
| CC | GO:0005739 | mitochondrion | 22 | 156 | 0.044943 |
| **(D) SAT lineage** | | | | | |
| BP | GO:0000003 | reproduction | 17 | 166 | 0.013897 |

| **(E) NIV lineage** | | | | | |
|---|---|---|---|---|---|
| BP | GO:0007610 | behavior | 1 | 1 | 8.22E-07 |
| BP | GO:0009835 | ripening | 1 | 1 | 7.09E-06 |
| CC | GO:0009536 | plastid | 159 | 526 | 3.86E-05 |
| CC | GO:0005576 | extracellular region | 15 | 36 | 0.000694 |
| MF | GO:0005102 | receptor binding | 2 | 3 | 0.006703 |
| MF | GO:0016740 | transferase activity | 84 | 269 | 0.007019 |
| MF | GO:0004872 | receptor activity | 4 | 8 | 0.009942 |
| BP | GO:0005975 | carbohydrate metabolic process | 46 | 141 | 0.024131 |
| BP | GO:0009908 | flower development | 28 | 92 | 0.028079 |
| BP | GO:0009991 | response to extracellular stimulus | 10 | 33 | 0.030031 |
| BP | GO:0009856 | pollination | 3 | 11 | 0.041472 |
| BP | GO:0000003 | reproduction | 39 | 166 | 0.043180 |

**Table S48. Summary of the detected PSGs orthologous to important functions in *O. rufipogon*, *O. sativa* and *O. nivara*.**

| Functional category | MSU gene model | Functional description | [a] Cloned orthologous genes |
|---|---|---|---|
| Flower development GO:0009908 | LOC_Os01g45760.1 | YUCCA-like gene | OsYUCCA1 |
| | LOC_Os02g02820.1 | tapetum degeneration retardation; basic helix-loop-helix protein | TDR |
| | LOC_Os02g02860.1 | glutamyl-tRNA synthetase | OsGluRS;Cde1(t) |
| | LOC_Os02g54600.1 | Mitogen activated protein Kinase Kinase 4;SMALL GRAIN 1 | OsMKK4;SMG1 |
| | LOC_Os03g05310.1 | Lesion mimic;Leaf senescence. | OsPAO |
| | LOC_Os03g12660.1 | Cytochrome P450 gene | OsDWARF4;CYP90B2 |
| | LOC_Os03g46190.1 | [b]QTL(Panicle length) | |
| | LOC_Os03g57660.1 | [b]QTL(Grain number) | |
| | LOC_Os03g59330.1 | [b]QTL(Grain number) | |
| | LOC_Os03g60430.2 | Oryza sativa INDETERMINATE SPIKELET 1 | OsIDS1 |
| | LOC_Os04g39470.1 | transcription factor with an MYB domain | OsMYB80 |
| | LOC_Os04g51000.1 | rice LFY homolog; ABERRANT PANICLE ORGANIZATION 2 | RFL;APO2 |
| | LOC_Os04g55560.2 | APETALA2 Transcription Factor | SHAT1 |
| | LOC_Os04g56170.1 | liguleless gene | OsLG1 |
| | LOC_Os06g04090.1 | NAC Transcription Factor | OsSWN1 |
| | LOC_Os06g16370.1 | Heading date 1 | Hd1 |
| | LOC_Os08g39390.1 | QTL(Stigma exsertion/awn length) | |
| | LOC_Os08g41940.1 | grain weight;squamosa promoter binding protein-like 16 | qGW8;OsSPL16 |
| | LOC_Os09g34070.1 | Spen-like protein | OsRRMh |
| | LOC_Os09g38790.1 | [b]QTL (Grain number) | |
| Ripening GO:0009835 | LOC_Os08g43654.1 | MATE efflux family protein, putative, expressed | |
| Reproduction GO:0000003 | LOC_Os01g45760.1 | YUCCA-like gene | OsYUCCA1 |
| | LOC_Os01g59660.1 | Gibberellin myb gene | OsGAmyb;MYBGA |
| | LOC_Os02g47970.1 | defective kernel 1 | OsDEK1 |
| | LOC_Os03g05310.1 | Lesion mimic;Leaf senescence. | OsPAO |
| | LOC_Os03g46190.1 | [b]QTL (Panicle length) | LOC_Os03g46190 |
| | LOC_Os03g49990.1 | slender rice 1; GRAS-domain protein | SLR1;OsGAI;Slr1-d |
| | LOC_Os03g58120.1 | [b]QTL (Grain number) | |
| | LOC_Os03g58540.1 | [b]QTL (Grain number) | |
| | LOC_Os03g59330.1 | [b]QTL (Grain number) | |
| | LOC_Os03g60430.2 | Oryza sativa INDETERMINATE SPIKELET 1 | OsIDS1 |
| | LOC_Os04g55560.2 | APETALA2 Transcription Factor | SHAT1 |
| | LOC_Os06g05090.1 | Protein arginine methyltransferase | OsPRMT10 |
| | LOC_Os06g41050.1 | COM1/SAE2 homolog | OsCOM1 |
| | LOC_Os06g45800.1 | [b]QTL (Grain length/grain weight) | |
| | LOC_Os09g26700.1 | [b]QTL (Grain weight) | |
| | LOC_Os09g34070.1 | Spen-like protein | OsRRMh |
| | LOC_Os09g35970.1 | [b]QTL (Grain number) | |
| | LOC_Os09g36450.1 | [b]QTL (Grain number) | |
| | LOC_Os10g28330.1 | early heading date 2; Cys-2/His-2-type zinc finger transcription factor | Ehd2;RID1;OsId1;Ghd10 |

| | | | |
|---|---|---|---|
| | LOC_Os01g45760.1 | YUCCA-like gene | OsYUCCA1 |
| | LOC_Os02g47970.1 | defective kernel 1 | OsDEK1 |
| | LOC_Os03g46190.1 | [b]QTL (Panicle length) | |
| | LOC_Os03g49990.1 | slender rice 1; GRAS-domain protein | SLR1;OsGAI;Slr1-d |
| | LOC_Os03g54100.1 | Two-Pore K+ Channel | OsTPKa |
| | LOC_Os03g57660.1 | [b]QTL (Grain number) | |
| | LOC_Os03g58120.1 | [b]QTL (Grain number) | |
| | LOC_Os03g58540.1 | [b]QTL (Grain number) | |
| Post-embryonic development GO:0009791 | LOC_Os03g60430.2 | Oryza sativa INDETERMINATE SPIKELET 1 | OsIDS1 |
| | LOC_Os03g63650.1 | BREVIS RADIX-like homologous gene in rice | OsBRXL2 |
| | LOC_Os04g55560.2 | APETALA2 Transcription Factor | SHAT1 |
| | LOC_Os06g05090.1 | Protein arginine methyltransferase | OsPRMT10 |
| | LOC_Os06g45800.1 | [b]QTL (Grain length/grain weight) | |
| | LOC_Os07g01810.1 | two pore potassium channel | OsTPKb |
| | LOC_Os09g26700.1 | [b]QTL (Grain weight) | |
| | LOC_Os09g34070.1 | Spen-like protein | OsRRMh |
| | LOC_Os09g35970.1 | [b]QTL (Grain number) | |
| | LOC_Os10g28330.1 | early heading date 2; Cys-2/His-2-type zinc finger transcription factor | Ehd2;RID1;OsId1;Ghd10 |
| Pollination GO:0009856 | LOC_Os08g39390 | [b]QTL (Stigma exsertion/awn length) | |
| | LOC_Os01g16810.1 | R2R3-type MYB gene;Carbon Starved Anther | CSA |
| | LOC_Os01g45760.1 | YUCCA-like gene | OsYUCCA1 |
| | LOC_Os01g59660.1 | Gibberellin myb gene | OsGAmyb;MYBGA |
| | LOC_Os01g66120.1 | NAC domain transcription factor | OsNAC6;SNAC2 |
| | LOC_Os02g01440.1 | nitric oxide association | OsNOA1 |
| | LOC_Os02g02820.1 | tapetum degeneration retardation; basic helix-loop-helix protein | TDR |
| | LOC_Os02g53690.1 | Growth Regulating Factor | OsGRF1;rhd1 |
| | LOC_Os02g54600.1 | Mitogen activated protein Kinase Kinase 4;SMALL GRAIN 1 | OsMKK4;SMG1 |
| | LOC_Os03g07140.1 | Fatty Acyl Carrier Protein Reductase; Defective Pollen Wall | DPW |
| | LOC_Os03g12660.1 | Cytochrome P450 gene | OsDWARF4;CYP90B2 |
| | LOC_Os03g21060.1 | plant-specific NAC transcriptional activator | OsNAP;PS1 |
| | LOC_Os03g49880.1 | rice TEOSINTE BRANCHED 1; TCP family transcription factor | OsTB1;FC1 |
| | LOC_Os03g51690.1 | KNOX family class 1 homeobox gene of rice | OSH1;Oskn1 |
| Multicellular organismal development GO:0007275 | LOC_Os03g53280.1 | [b]QTL (Grain number) | LOC_Os03g53280 |
| | LOC_Os03g57660.1 | [b]QTL (Grain number) | LOC_Os03g57660 |
| | LOC_Os03g58910.2 | [b]QTL (Grain number) | LOC_Os03g58910 |
| | LOC_Os03g60430.2 | Oryza sativa INDETERMINATE SPIKELET 1 | OsIDS1 |
| | LOC_Os03g60650.1 | [b]QTL (Grain number) | LOC_Os03g60650 |
| | LOC_Os03g63650.1 | BREVIS RADIX-like homologous gene in rice | OsBRXL2 |
| | LOC_Os04g39470.1 | transcription factor with an MYB domain | OsMYB80 |
| | LOC_Os04g46470.1 | carotenoid cleavage dioxygenase;high tillering dwarf 1; semidwarf-t;dwarf and increased tillering1 | OsCCD7;htd1;sd-t;dit1 |
| | LOC_Os04g51000.1 | rice LFY homolog; ABERRANT PANICLE ORGANIZATION 2 | RFL;APO2 |
| | LOC_Os04g55560.2 | APETALA2 Transcription Factor | SHAT1 |
| | LOC_Os04g56170.1 | liguleless gene | OsLG1 |
| | LOC_Os05g03884.1 | KNOX family class 1 homeobox gene of rice | OSH71;Oskn2 |
| | LOC_Os06g04090.1 | NAC Transcription Factor | OsSWN1 |
| | LOC_Os06g06050.1 | bungetsuwaito tillering dwarf; dwarf-3 | D3;SOLS |
| | LOC_Os06g10990.1 | Hybrid sterility | ORF3 |
| | LOC_Os06g40550.1 | ABC Transporter Protein; Post-meiotic Deficicent Anther1 | OsABCG15;PDA1 |

| | | | |
|---|---|---|---|
| | LOC_Os07g03770.1 | ebisumochi dwarf; dwarf 6;<br>KNOX family class 1 homeobox gene of rice | d6;OSH15;Oskn3 |
| | LOC_Os08g29660.1 | WRKY transcription factor | OsWRKY69 |
| | LOC_Os08g39390.1 | [b]QTL (Stigma exsertion/awn length) | LOC_Os08g39390 |
| | LOC_Os08g39390.1 | [b]QTL (Stigma exsertion/awn length) | LOC_Os08g39390 |
| | LOC_Os09g34070.1 | Spen-like protein | OsRRMh |
| | LOC_Os09g36450.1 | [b]QTL (Grain number) | LOC_Os09g36450 |
| | LOC_Os10g28330.1 | early heading date 2;<br>Cys-2/His-2-type zinc finger transcription factor | Ehd2;RID1;OsId1;Ghd10 |
| | LOC_Os10g39800.1 | [b]QTL (Grain width) | LOC_Os10g39800 |
| | LOC_Os11g08210.1 | NAC domain transcription factor | OsNAC5 |
| Response to extracellular stimulus GO:0009991 | LOC_Os02g52210.1 | [b]QTL (Grain length) | |
| | LOC_Os05g48390.1 | ubiquitin-conjugating E2 enzyme; leaf tip necrosis1 | LTN1;OsPHO2;OsUBC35 |
| | LOC_Os07g48790.1 | [b]QTL (Awn length/heading date) | |
| | LOC_Os10g36924.1 | boric acid channel; Dwarf and tiller-enhancing 1 | OsNIP3;1;DTE1 |
| | LOC_Os10g42960.1 | high-affinity urea transporter | OsDUR3 |
| Response to biotic stimulus GO:0009607 | LOC_Os01g43550.1 | WRKY transcription factor | OsWRKY12;OsWRKY03 |
| | LOC_Os02g08440.1 | WRKY transcription factor | OsWRKY71 |
| | LOC_Os02g53180.1 | aminocyclopropane-1-carboxylate oxidase gene | OsACO3 |
| | LOC_Os02g54600.1 | Mitogen activated protein Kinase Kinase 4;SMALL GRAIN 1 | OsMKK4;SMG1 |
| | LOC_Os03g05310.1 | Lesion mimic;Leaf senescence. | OsPAO |
| | LOC_Os03g60960.1 | [b]QTL (Grain number) | LOC_Os03g60960 |
| | LOC_Os04g54474.1 | bZIP transcription factor;<br>Oryza sativa TGA factor for phytoalexin | OsTGAP1;OsbZIP37 |
| | LOC_Os06g11010.1 | wide compatibility gene;Hybrid sterility-5 | S5 |
| | LOC_Os08g04500.1 | (E)-β-caryophyllene synthase（抗虫性） | OsTPS3 |
| | LOC_Os08g29660.1 | WRKY transcription factor | OsWRKY69 |
| | LOC_Os08g37874.1 | [b]QTL (Stigma exsertion/awn length) | LOC_Os08g37874 |
| | LOC_Os09g35970.1 | [b]QTL (Grain number) | LOC_Os09g35970 |
| Photosynthesis GO:0015979 | LOC_Os01g16040.1 | 3′-Phosphoadenosine 5′-Phosphosulfate Transporter1 | PAPST1 |
| | LOC_Os03g56869.1 | [b]QTL (Grain number) | LOC_Os03g56869 |
| | LOC_Os09g36450.1 | [b]QTL (Grain number) | LOC_Os09g36450 |
| Embryo development GO:0009790 | LOC_Os01g45760.1 | YUCCA-like gene | OsYUCCA1 |
| | LOC_Os01g68370.1 | B3 domain-containing protein | OSVP1;VP1 |
| | LOC_Os02g47970.1 | defective kernel 1 | OsDEK1 |
| | LOC_Os03g57660.1 | [b]QTL (Grain number) | |
| | LOC_Os03g58120.1 | [b]QTL (Grain number) | |
| | LOC_Os03g58540.1 | [b]QTL (Grain number) | |
| | LOC_Os03g58910.2 | [b]QTL (Grain number) | |
| | LOC_Os06g45800.1 | [b]QTL (Grain length/grain weight) | |
| | LOC_Os09g26700.1 | [b]QTL (Grain weight) | |
| | LOC_Os09g34070.1 | Spen-like protein | OsRRMh |
| | LOC_Os09g35970.1 | [b]QTL (Grain number) | |
| Response to stress GO:0006950 | LOC_Os01g39630.1 | rice homolog of human RAD51C | RAD51C;OsRAD51C |
| | LOC_Os01g40670.1 | telomere repeat-binding factor | OsTRBF1 |
| | LOC_Os01g43550.1 | WRKY transcription factor | OsWRKY12;OsWRKY03 |
| | LOC_Os01g51154.1 | telomere repeat-binding factor | OsTRBF3 |
| | LOC_Os01g57210.1 | katanin P80 ortholog;microtubule-severing enzyme | OsKTN80c |

| | | |
|---|---|---|
| LOC_Os01g66120.1 | NAC domain transcription factor | OsNAC6;SNAC2 |
| LOC_Os02g01440.1 | nitric oxide association | OsNOA1 |
| LOC_Os02g08440.1 | WRKY transcription factor | OsWRKY71 |
| LOC_Os02g51670.1 | [b]QTL (Grain length) | |
| LOC_Os02g52460.1 | [b]QTL (Grain length) | |
| LOC_Os02g52780.1 | bZIP transcription factor | OsbZIP23 |
| LOC_Os02g54600.1 | Mitogen activated protein Kinase Kinase 4; SMALL GRAIN 1 | OsMKK4;SMG1 |
| LOC_Os02g58590.1 | *N-acetylglucosaminyltransferase I* | GnT1 |
| LOC_Os03g05310.1 | Lesion mimic;Leaf senescence. | OsPAO |
| LOC_Os03g08460.1 | EREBP transcription factor | OsEBP-89 |
| LOC_Os03g12414.1 | cyclin; SOLO DANCERS | SDS;OsSDS |
| LOC_Os03g45330.1 | [b]QTL (Panicle length) | |
| LOC_Os03g49990.1 | slender rice 1; GRAS-domain protein | SLR1;OsGAI;Slr1-d |
| LOC_Os03g50210.1 | [b]QTL (Grain number) | |
| LOC_Os03g51610.1 | inositol 1,3,4-trisphosphate 5/6-kinase gene | OsITPK3 |
| LOC_Os03g51920.1 | [b]QTL (Grain number) | |
| LOC_Os03g53720.1 | [b]QTL (Grain number) | |
| LOC_Os03g58390.1 | [b]QTL (Grain number) | |
| LOC_Os03g60960.1 | [b]QTL (Grain number) | |
| LOC_Os04g54474.1 | bZIP transcription factor; Oryza sativa TGA factor for phytoalexin | OsTGAP1;OsbZIP37 |
| LOC_Os05g09050.1 | [b]QTL (Grain width) | |
| LOC_Os05g27930.1 | AP2/EREBP transcription factor gene | OsDREB2B |
| LOC_Os05g41880.1 | MutS-homolog gene | OsMSH5 |
| LOC_Os05g48390.1 | ubiquitin-conjugating E2 enzyme; leaf tip necrosis1 | LTN1;OsPHO2;OsUBC35 |
| LOC_Os06g06050.1 | bungetsuwaito tillering dwarf; dwarf-3 | D3;SOLS |
| LOC_Os06g10990.1 | Hybrid sterility | ORF3 |
| LOC_Os06g11010.1 | wide compatibility gene;Hybrid sterility-5 | S5 |
| LOC_Os06g41050.1 | COM1/SAE2 homolog | OsCOM1 |
| LOC_Os07g09060.1 | Rice Aldehyde Dehydrogenase | OsALDH6B2 |
| LOC_Os07g34570.1 | defense responsive gene 8; putative thiamine synthase | OsDR8;OsXNP |
| LOC_Os07g39480.1 | WRKY transcription factor 78 | OsWRKY78 |
| LOC_Os07g46410.1 | rice chloroplast NADPH thioredoxin reductase | NTRC |
| LOC_Os07g48870.1 | [b]QTL (Awn length/heading date) | |
| LOC_Os08g29660.1 | WRKY transcription factor | OsWRKY69 |
| LOC_Os09g28310.1 | bZIP transcription factor | OsbZIP72 |
| LOC_Os09g35970.1 | [b]QTL (Grain number) | |
| LOC_Os10g41200.1 | Rice MYB | MYBS3 |
| LOC_Os10g42960.1 | high-affinity urea transporter | OsDUR3 |
| LOC_Os11g08210.1 | NAC domain transcription factor | OsNAC5 |

[a] These orthologous cloned gene were retrieved from the database CHINA RICE DATA CENTER (http://www.ricedata.cn/);

[b] These genes were found in quantitative trait loci (QTLs) associated with domestication traits identified previously (Huang et al. 2012).
